# Coronary atherosclerosis: Un-altered physiology

**DOI:** 10.1101/581454

**Authors:** Paul de Groot, Roel W Veldhuizen

## Abstract

**Background:** End-stage coronary artery atherosclerosis has been studied extensively but the exact mechanisms of initiation and progression have not been defined fully. The aim of this study was to mathematically describe luminal change in relation to coronary vessel wall thickness in its progression from normal to atherosclerotic to establish whether these explain the pathophysiology.

**Methods:** One hundred coronary artery sections were graded histologically as ‘normal’ to ‘highly atherosclerotic’. Random systemic sampling by image analysis yielded 32 measurements (lumen radius and intima, medial, and adventitial thickness) from each section along 32 evenly spaced radii.

**Results:** The raw data follow an undulating course in relation to successive segments in all sections analyzed, pointing to a dynamic and well-ordered system. The calculated values, studied in triplets, followed a non-synchronized parabolic course, which was converted to linearity by taking the change in numbers (n-(n-1)=Δ) into account. The course and sign of ‘Δvessel wall’ (resulting from summed Δintima, Δmedia, Δadventitia) and ‘Δlumen radius’ values were unique for each triplet. Triplets order according to ‘Δlumen radius minus Δvessel wall’ and its course given by the trendline a-value presented stages in which increased Δvessel wall resulted in increased Δlumen radius in stages 1 and 3 and decreased Δlumen radius in stage 2. This phenomenon was found in all sections regardless of histological indication and independent of vessel wall constituent parts (intima, media, adventitia).

**Conclusions:** Similar basic processes are defined in all sections regardless of histological rating, indicating un-altered physiology. As such, coronary atherosclerosis can only be defined by a large to small shift of the triplets Δvessel wall trendline a-value. Consequently, no parameter of vessel wall pathology exists in absolute terms. Vessel wall composition has no importance for Δlumen radius.

## Introduction

The anatomy and pathology of the coronary arteries are widely studied, especially by histological means. However, basic knowledge is lacking regarding, for example, the mechanism of arterial tapering, which corresponds to deficient knowledge of vessel wall/lumen interactions and, even more fundamentally, the intimal, medial, adventitial interplay.

Research has traditionally focused on the histopathology of plaque formation, the end-stage atherosclerotic process. The process from initiation to plaque formation has often been studied in a retrograde manner [1–3], which may lead to cause/effect reversal. An enlarged intima was considered a characteristic and regarded as a key player in atherosclerotic pathology [4, 5]. Consequently, the limits of normality, the thickness of the media and adventitia, and their role in atherosclerosis remain elusive.

We hypothesized that mathematical differences in the cross-sectional dimensions of normal and atherosclerotic arteries between stages could be used to help explain the pathophysiology of coronary artery atherosclerosis. This study aims to bridge the knowledge gap between the physiology of vessel morphology and the pathogenesis of atherosclerosis by mathematically defining both processes. Using a specially designed morphometric method termed ‘random systemic sampling’ [6], we aim to define how vessel wall thickness and lumen size are related and how the vessel wall layers (intima, media, adventitia) contribute to the pathophysiology of atherosclerosis.

## Material and Methods

### Study population and ethical approval

Study samples were obtained from autopsies of unselected patients regardless of the cause of death. No inclusion or exclusion criteria were applied. A total 100 cases were accepted and included for study. The Haaglanden MC ethics committee approved the study protocol and the use of specimens obtained from routine post-mortem examinations.

### Coronary artery sampling and histology

Tissue blocks from the heart with coronary vessels attached were fixed by submersion in 4% phosphate-buffered formaldehyde solution (0.1M, pH 7.0) for 48 hours prior to further sampling. The arteries were not pressure perfused in order to retain ‘natural’ residual vascular wall stress. Vessel cross-sections were selected at random with no preference for left or right coronary arteries, sampling site, or cause of death. Sections were processed using routine methods, paraffin embedded, and 5-μm sections cut and stained with hematoxylin and eosin for microscopic assessment. Morphometric assessment was preferentially performed on von Gieson/elastica stained sections.

### Dataset

All sections were imaged and assessed by microscopy. Sections were histologically defined as follows: normal (intima < media), circumferential non-atherosclerotic enlarged vessel wall (intima ≥ media), circumferential and unilateral atherosclerotic enlarged vessel wall (signs of atherosclerosis, i.e., pattern of acellularity, cholesterol deposition, calcification, fibrosis, inflammation). Thus, all sections together represented the course from ‘normal’ (no signs of atherosclerosis) to ‘circumferential atherosclerosis’. One fully analyzed specimen from each histologically defined group highlighted the mathematical changes during progression. An additional section was histologically rated as ‘extreme unilateral atherosclerosis’ (not shown).

### Image analysis

Computer-assisted image analysis was performed using ZEN 2 lite software (Carl Zeiss). Digitized tissue sections were displayed on a monitor and the center of each vessel located using a computerized best fit circle procedure. Only sections with an external elastic lamina Feret circle of at least 0.8 were used to exclude sectioning artefacts and interpretation bias.

Using a random starting position, 32 radii were drawn from the center (i.e., one every 11.25 degrees) (random *systemic* sampling; Fig 1). The observer then identified and marked the positions of the endothelium, internal and external elastic lamina, and the adventitial-fat border along each radius. The lumen radius (Rlu) and the dimensions of the intima, media, and adventitia (INTd, MEDd, and ADVd, respectively) at each location were calculated in micrometers by the software. Matched values along each radius were jointly termed the ‘measurement unit’. Thirty-two measurement units were available for each section. All data were tabulated for further analysis.

**Fig 1.**
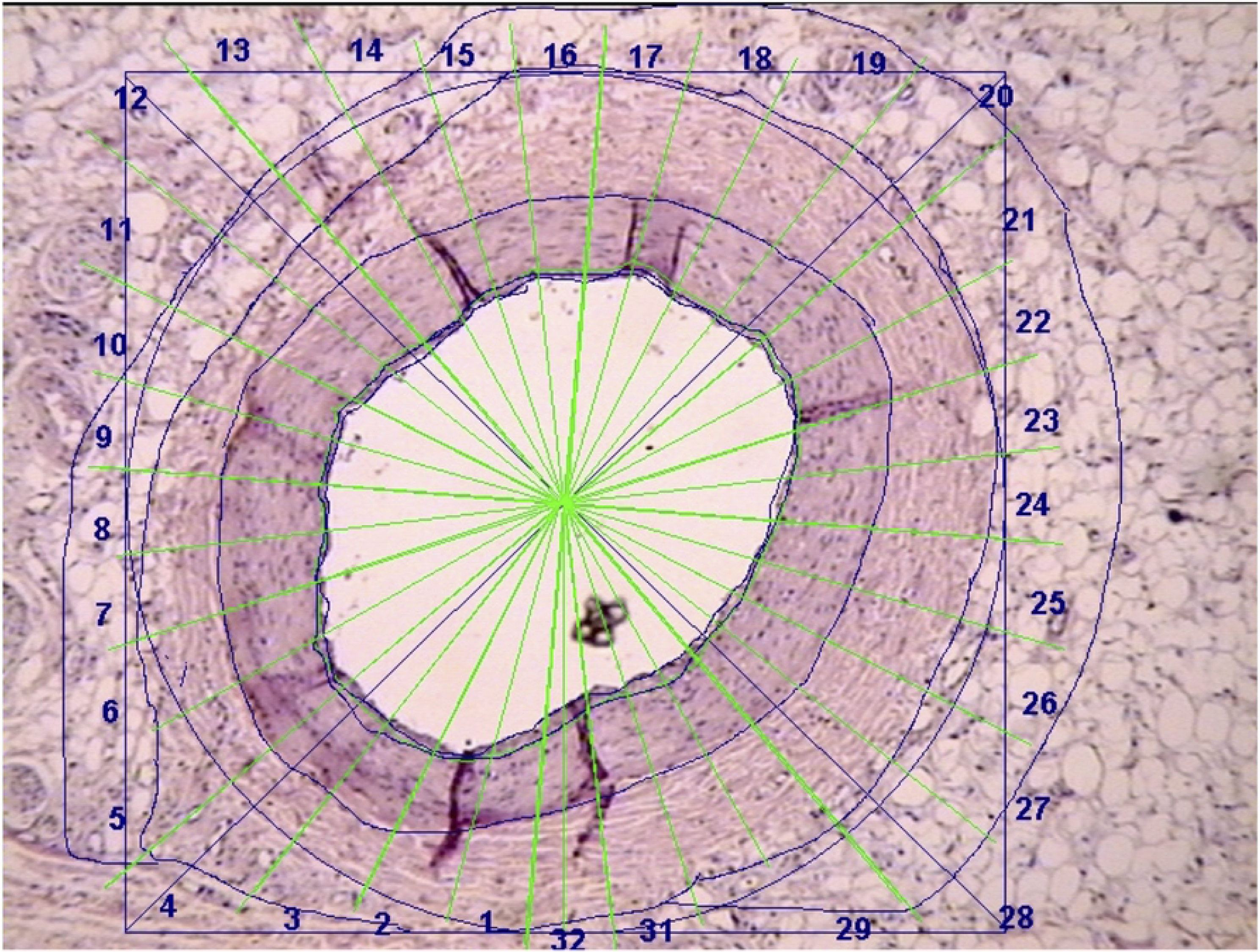
Random systemic sampling computer-assisted image analysis. The center point is located by the best fit circle procedure. Thirty-two radii differing by 11.25 degrees defined 32 measurement units of the lumen radius and intima, media, and adventitia dimensions.

### Validation of the term value as a computerized classification parameter

INTd, MEDd, and ADVd were highly variable, even between two consecutive measurement units and regardless of the originating section. As the three vessel wall layer dimensions were not usually independent, they were defined as a single functional unit (FU-1). Similarly, Rlu-Wd, VWd, and Rlu defined a functional unit (FU-2). The relationship between the three values constituting FU-1 and FU-2 was parabolic and interrelated according to the formula ax^2^ + bx + c. A constant characteristic of each functional unit (the ‘term value’; T-FU1 and T-FU2) was produced by subtracting the difference between ADVd and MEDd from the difference between MEDd and INTd, and subtracting Rlu-VWd from VWd-(Rlu-VWd), respectively.

### Process study

Data from random systemic sampling represented a *static* state, a snapshot in time. Each segment provided FU-1 and FU-2, which were characterized by T-FU1 and T-FU2, respectively. T-FU2 values set against consecutive segment numbers showed a well-ordered saw-toothed course in each section; the upward course changed to a downward course at an acute angle (Fig 2–5), indicating a *dynamic* process.

**Fig 2.**
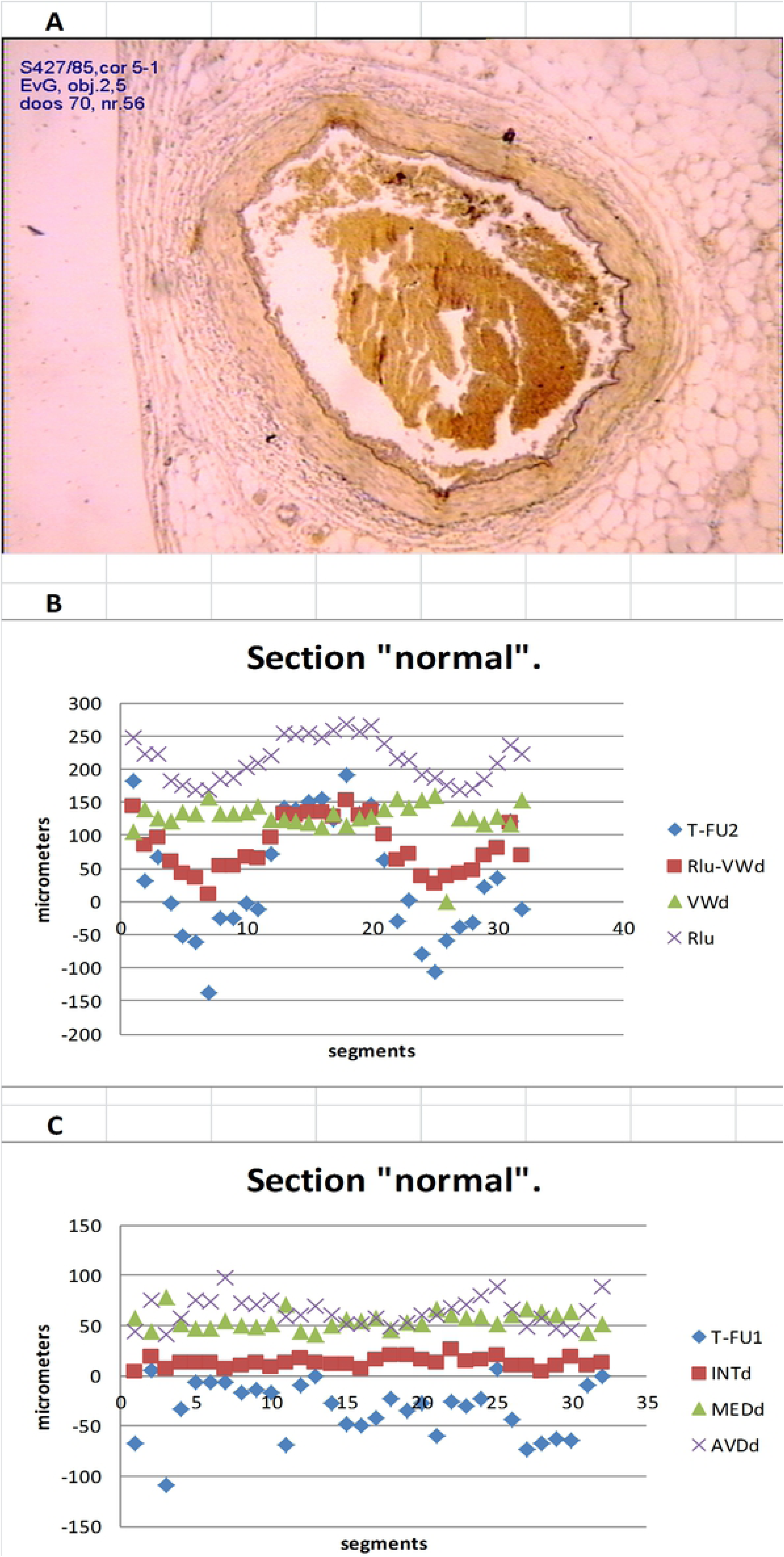
Section rated ‘normal’. (A) Micrograph. (B) Scatter diagram T-FU2, Rlu-VWd, VWd, and Rlu vs. segment numbers. (C) T-FU1, INTd, MEDd, and ADVd vs. segment numbers.

The changes in the INTd, MEDd, and ADVd and Rlu-VWd, VWd, and Rlu of consecutive segments were studied in triplets. Each triplet provided three successive values of FU-1 and FU-2 defined by their collective parabolic course and represented by their corresponding trendline formula (R^2^ =1). Intermediate segment values were calculated by interpolation. For example, triplet (1-3): segment numbers (x) and segment numbers squares (x^2^) interpolated in formula y=ax^2^ + bx + c provided calculated y values of x= 1; 1.1; 1.2; … 2; 2.1; … 3. Therefore, each section provided 16 triplets (1-3), (3-5), … (31-1), each consisting of 20 calculated FU-1 and FU-2 component values. The non-synchronized parabolic course of INTd, MEDd, and ADVd and Rlu-VWd, VWd, and Rlu in relation to T-FU1 and T-FU2, respectively, were marked by strong variability in peak/bottom parabola values, which made mutual comparisons impossible. Therefore, a step by step course was described by the value change (Δvalues 1.1-1.0 … n-(n-1)), which transformed the parabolic relationship into a linear one (trendline formula: y = ax + b, R^2^=1).

## Results

### Measurement data

Measurement data for ‘normal’ (Table 1, Fig 2), ‘circumferentially enlarged intima’ (Table 2, Fig 3), ‘unilateral atherosclerosis’ (Table 3, Fig 4), and ‘circumferential atherosclerosis’ sections (Table 4, Fig 5) showed the saw-toothed course of T-FU2 in relation to segment number. The additional ‘extreme unilateral atherosclerosis’ section (not shown) also presented this phenomenon. T-FU2 curves were similar to Rlu-VWd curves, and to a lesser extent Rlu curves. The VWd and Rlu were generally antagonistic, but some segments followed a congruent course. An increase in the VWd was not always accompanied by a decrease in Rlu, and vice versa. The INTd, MEDd, and ADVd also alternated, but did not follow a distinct pattern.

**Table 1.** Measurement Values of 32 Segments in a ‘Normal’ Section.

| Segment | **T-FU2** | Rlu-VWd | VWd | Rlu | **T-FU1** | INTd | MEDd | AVDd |
| --- | --- | --- | --- | --- | --- | --- | --- | --- |
| 1 | **181** | 143 | 105 | 248 | **-68** | 3 | 58 | 45 |
| 2 | **32** | 85 | 138 | 223 | **5** | 18 | 44 | 76 |
| 3 | **68** | 97 | 126 | 223 | **-109** | 7 | 79 | 41 |
| 4 | **-2** | 59 | 121 | 181 | **-34** | 13 | 52 | 57 |
| 5 | **-52** | 41 | 135 | 176 | **-6** | 13 | 47 | 75 |
| 6 | **-62** | 36 | 133 | 168 | **-7** | 13 | 47 | 74 |
| 7 | **-138** | 10 | 158 | 168 | **-6** | 6 | 55 | 97 |
| 8 | **-25** | 54 | 132 | 185 | **-17** | 10 | 50 | 72 |
| 9 | **-25** | 54 | 133 | 186 | **-14** | 13 | 49 | 71 |
| 10 | **-2** | 67 | 136 | 203 | **-17** | 9 | 51 | 76 |
| 11 | **-11** | 66 | 143 | 208 | **-69** | 13 | 70 | 59 |
| 12 | **72** | 97 | 123 | 220 | **-9** | 18 | 44 | 61 |
| 13 | **141** | 132 | 122 | 254 | **0** | 13 | 41 | 69 |
| 14 | **139** | 130 | 121 | 251 | **-28** | 11 | 50 | 60 |
| 15 | **150** | 134 | 119 | 253 | **-48** | 11 | 56 | 52 |
| 16 | **155** | 134 | 113 | 247 | **-49** | 7 | 54 | 52 |
| 17 | **124** | 128 | 131 | 259 | **-42** | 16 | 58 | 58 |
| 18 | **190** | 152 | 115 | 267 | **-22** | 20 | 46 | 49 |
| 19 | **133** | 129 | 126 | 256 | **-35** | 20 | 54 | 53 |
| 20 | **147** | 137 | 128 | 265 | **-27** | 16 | 52 | 60 |
| 21 | **62** | 100 | 138 | 238 | **-60** | 13 | 66 | 60 |
| 22 | **-30** | 62 | 154 | 216 | **-25** | 27 | 60 | 68 |
| 23 | **2** | 72 | 142 | 214 | **-31** | 14 | 58 | 70 |
| 24 | **-80** | 37 | 154 | 190 | **-23** | 16 | 59 | 79 |
| 25 | **-107** | 26 | 160 | 186 | **7** | 20 | 51 | 89 |
| 26 | **-59** | 39 | 136 | 175 | **-44** | 10 | 60 | 66 |
| 27 | **-39** | 43 | 125 | 168 | **-74** | 10 | 66 | 49 |
| 28 | **-31** | 47 | 125 | 172 | **-67** | 3 | 64 | 58 |
| 29 | **22** | 69 | 116 | 185 | **-63** | 9 | 60 | 47 |
| 30 | **35** | 81 | 128 | 209 | **-64** | 19 | 64 | 45 |
| 31 | **121** | 119 | 116 | 235 | **-10** | 10 | 42 | 65 |
| 32 | **-13** | 70 | 153 | 223 | **0** | 13 | 51 | 89 |

Segment clusters comprise 1-7, 8-17, 18-28, and 19-32.

**Table 2.** Measurement Values of 32 Segments in a ‘Circumferential Enlarged Intima’ Section.

| Segment | T-FU2 | r-lu-VW | VW-dik | r-lumen | T-FU1 | intdik | meddik | advdik |
| --- | --- | --- | --- | --- | --- | --- | --- | --- |
| 1 | **-608** | 53 | 713 | 766 | **194** | 252 | 173 | 288 |
| 2 | **-462** | 114 | 690 | 804 | **116** | 244 | 191 | 254 |
| 3 | **-349** | 171 | 690 | 861 | **92** | 230 | 200 | 261 |
| 4 | **-370** | 180 | 730 | 910 | **96** | 238 | 211 | 280 |
| 5 | **-171** | 270 | 711 | 981 | **67** | 226 | 215 | 270 |
| 6 | **29** | 363 | 697 | 1060 | **128** | 223 | 190 | 284 |
| 7 | **157** | 426 | 696 | 1123 | **57** | 204 | 213 | 280 |
| 8 | **95** | 410 | 725 | 1135 | **235** | 268 | 163 | 294 |
| 9 | **407** | 523 | 638 | 1161 | **74** | 225 | 188 | 225 |
| 10 | **657** | 620 | 583 | 1203 | **96** | 211 | 162 | 210 |
| 11 | **701** | 631 | 562 | 1193 | **55** | 166 | 169 | 227 |
| 12 | **435** | 515 | 594 | 1109 | **94** | 160 | 167 | 268 |
| 13 | **368** | 461 | 554 | 1015 | **41** | 172 | 171 | 211 |
| 14 | **58** | 335 | 611 | 946 | **65** | 164 | 182 | 265 |
| 15 | **114** | 338 | 562 | 899 | **3** | 172 | 186 | 204 |
| 16 | **-101** | 249 | 599 | 847 | **45** | 197 | 185 | 217 |
| 17 | **-342** | 155 | 652 | 808 | **243** | 300 | 136 | 216 |
| 18 | **-491** | 109 | 708 | 817 | **345** | 317 | 121 | 270 |
| 19 | **-380** | 156 | 693 | 849 | **299** | 280 | 131 | 281 |
| 20 | **-563** | 103 | 769 | 872 | **250** | 297 | 173 | 300 |
| 21 | **-529** | 136 | 801 | 937 | **332** | 293 | 156 | 352 |
| 22 | **-489** | 172 | 833 | 1005 | **376** | 272 | 152 | 408 |
| 23 | **63** | 377 | 691 | 1068 | **355** | 287 | 112 | 292 |
| 24 | **452** | 544 | 637 | 1181 | **298** | 268 | 113 | 256 |
| 25 | **978** | 750 | 521 | 1270 | **79** | 194 | 147 | 180 |
| 26 | **718** | 656 | 594 | 1250 | **160** | 252 | 145 | 197 |
| 27 | **433** | 538 | 643 | 1181 | **316** | 245 | 109 | 289 |
| 28 | **315** | 463 | 611 | 1074 | **243** | 237 | 123 | 252 |
| 29 | **70** | 358 | 646 | 1004 | **301** | 227 | 115 | 303 |
| 30 | **-126** | 258 | 643 | 901 | **219** | 227 | 141 | 274 |
| 31 | **-357** | 158 | 673 | 830 | **222** | 235 | 150 | 288 |
| 32 | **-554** | 80 | 714 | 794 | **222** | 235 | 164 | 315 |

Segment clusters comprise 1-10, 11-14, 15-18, 19-22, 23-25, and 26-32.

**Table 3.** Measurement Values of 32 Segments in a ‘Unilateral Atherosclerosis’ Section.

| Segment | T-FU2 | rlu-VW | VWdik | Rlu | T-FU1 | INTd | MEDd | ADVd |
| --- | --- | --- | --- | --- | --- | --- | --- | --- |
| 1 | **-1528** | -469 | 590 | 121 | **483** | 433 | 36 | 121 |
| 2 | **-1608** | -498 | 611 | 112 | **519** | 464 | 31 | 116 |
| 3 | **-1741** | -543 | 655 | 113 | **576** | 493 | 26 | 136 |
| 4 | **-1823** | -569 | 685 | 116 | **545** | 499 | 47 | 140 |
| 5 | **-1671** | -509 | 653 | 145 | **477** | 497 | 59 | 98 |
| 6 | **-913** | -201 | 511 | 311 | **334** | 357 | 59 | 96 |
| 7 | **-356** | 28 | 412 | 440 | **219** | 240 | 64 | 107 |
| 8 | **-199** | 100 | 399 | 499 | **245** | 207 | 51 | 141 |
| 9 | **260** | 286 | 312 | 598 | **139** | 118 | 58 | 136 |
| 10 | **215** | 263 | 311 | 574 | **172** | 92 | 46 | 172 |
| 11 | **156** | 231 | 306 | 537 | **146** | 83 | 54 | 169 |
| 12 | **163** | 228 | 292 | 520 | **119** | 61 | 58 | 173 |
| 13 | **310** | 272 | 234 | 506 | **63** | 55 | 57 | 122 |
| 14 | **309** | 276 | 244 | 520 | **94** | 48 | 50 | 147 |
| 15 | **462** | 328 | 194 | 522 | **48** | 44 | 49 | 101 |
| 16 | **220** | 234 | 247 | 481 | **119** | 73 | 43 | 131 |
| 17 | **207** | 227 | 247 | 474 | **58** | 76 | 63 | 108 |
| 18 | **141** | 202 | 263 | 465 | **76** | 81 | 62 | 119 |
| 19 | **91** | 180 | 270 | 450 | **62** | 75 | 69 | 126 |
| 20 | **4** | 145 | 286 | 432 | **37** | 77 | 83 | 127 |
| 21 | **-1** | 146 | 292 | 438 | **73** | 78 | 73 | 141 |
| 22 | **112** | 195 | 277 | 472 | **93** | 86 | 61 | 130 |
| 23 | **183** | 231 | 280 | 511 | **113** | 68 | 56 | 157 |
| 24 | **192** | 246 | 301 | 547 | **169** | 81 | 44 | 177 |
| 25 | **59** | 208 | 358 | 566 | **169** | 71 | 63 | 224 |
| 26 | **-113** | 126 | 365 | 491 | **119** | 113 | 82 | 170 |
| 27 | **-449** | -11 | 427 | 415 | **207** | 170 | 73 | 184 |
| 28 | **-383** | -1 | 381 | 380 | **212** | 189 | 56 | 135 |
| 29 | **-578** | -87 | 404 | 317 | **234** | 246 | 57 | 102 |
| 30 | **-850** | -200 | 450 | 249 | **314** | 311 | 45 | 93 |
| 31 | **-1224** | -347 | 529 | 182 | **428** | 368 | 34 | 127 |
| 32 | **-1286** | -381 | 524 | 143 | **432** | 391 | 31 | 102 |

Segment clusters comprise 1-4, 5-8, 9-11, 12-15, 16-21, 22-24, and 25-32.

**Table 4.** Measurement Values of 32 Segments in a ‘Circumferential Atherosclerosis’ Section.

| Segment | T-FU2 | rlu-VW | VWdik | Rlu | T-FU1 | INTd | MEDd | ADVd |
| --- | --- | --- | --- | --- | --- | --- | --- | --- |
| 1 | **-1972** | -493 | 986 | 493 | **718** | 724 | 89 | 173 |
| 2 | **-2201** | -573 | 1055 | 482 | **639** | 718 | 139 | 198 |
| 3 | **-2315** | -609 | 1097 | 488 | **634** | 707 | 154 | 236 |
| 4 | **-2470** | -654 | 1162 | 507 | **665** | 764 | 166 | 232 |
| 5 | **-2578** | -674 | 1230 | 555 | **763** | 785 | 156 | 289 |
| 6 | **-2612** | -668 | 1275 | 607 | **818** | 780 | 153 | 343 |
| 7 | **-2351** | -561 | 1228 | 666 | **791** | 774 | 146 | 308 |
| 8 | **-2081** | -448 | 1185 | 736 | **744** | 760 | 147 | 278 |
| 9 | **-1796** | -325 | 1146 | 821 | **738** | 721 | 136 | 288 |
| 10 | **-238** | 286 | 810 | 1096 | **376** | 383 | 145 | 283 |
| 11 | **-466** | 185 | 837 | 1022 | **484** | 450 | 118 | 269 |
| 12 | **-911** | 2 | 916 | 919 | **542** | 472 | 125 | 319 |
| 13 | **-1585** | -283 | 1020 | 737 | **693** | 600 | 109 | 311 |
| 14 | **-1765** | -377 | 1012 | 635 | **545** | 575 | 156 | 281 |
| 15 | **-2143** | -543 | 1057 | 514 | **654** | 657 | 134 | 266 |
| 16 | **-2293** | -614 | 1065 | 450 | **503** | 665 | 187 | 213 |
| 17 | **-2464** | -682 | 1101 | 420 | **645** | 739 | 152 | 210 |
| 18 | **-2713** | -774 | 1164 | 390 | **732** | 796 | 144 | 224 |
| 19 | **-3056** | -893 | 1271 | 378 | **908** | 836 | 121 | 314 |
| 20 | **-3553** | -1057 | 1438 | 381 | **941** | 882 | 166 | 390 |
| 21 | **-3661** | -1086 | 1489 | 404 | **1044** | 907 | 148 | 434 |
| 22 | **-3704** | -1080 | 1544 | 464 | **1092** | 923 | 151 | 470 |
| 23 | **-3310** | -926 | 1459 | 533 | **1065** | 861 | 131 | 467 |
| 24 | **-2149** | -480 | 1189 | 709 | **865** | 702 | 108 | 378 |
| 25 | **-1936** | -393 | 1151 | 758 | **916** | 742 | 78 | 330 |
| 26 | -1108 | -65 | 978 | 913 | **771** | 591 | 69 | 318 |
| 27 | -1008 | -38 | 931 | 893 | **713** | 573 | 73 | 285 |
| 28 | **-1517** | -269 | 978 | 708 | **837** | 667 | 47 | 264 |
| 29 | **-1450** | -257 | 935 | 678 | **734** | 604 | 67 | 264 |
| 30 | **-1539** | -304 | 930 | 626 | **733** | 629 | 66 | 235 |
| 31 | **-1694** | -374 | 946 | 571 | **677** | 657 | 90 | 199 |
| 32 | **-1935** | -467 | 1002 | 535 | **723** | 708 | 93 | 201 |

Segment clusters comprise 1-6, 7-9, 10-19, 20-27, and 28-32.

**Fig 3.**
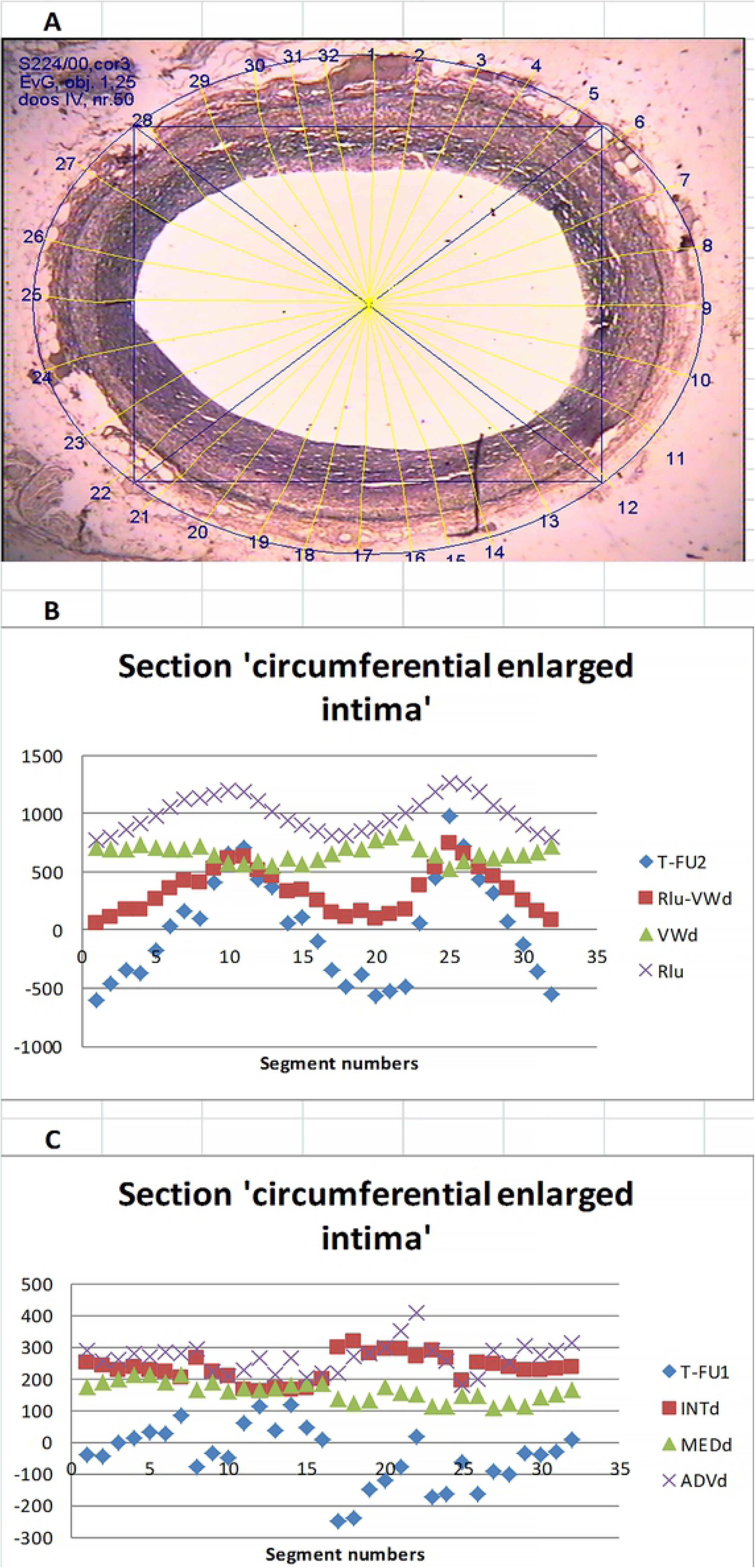
Section rated ‘circumferential enlarged intima’. (A) Micrograph. (B) Scatter diagram T-FU2, Rlu-VWd, VWd, and Rlu vs. segment numbers. (C) T-FU1, INTd, MEDd, and ADVd vs. segment numbers.

**Fig 4.**
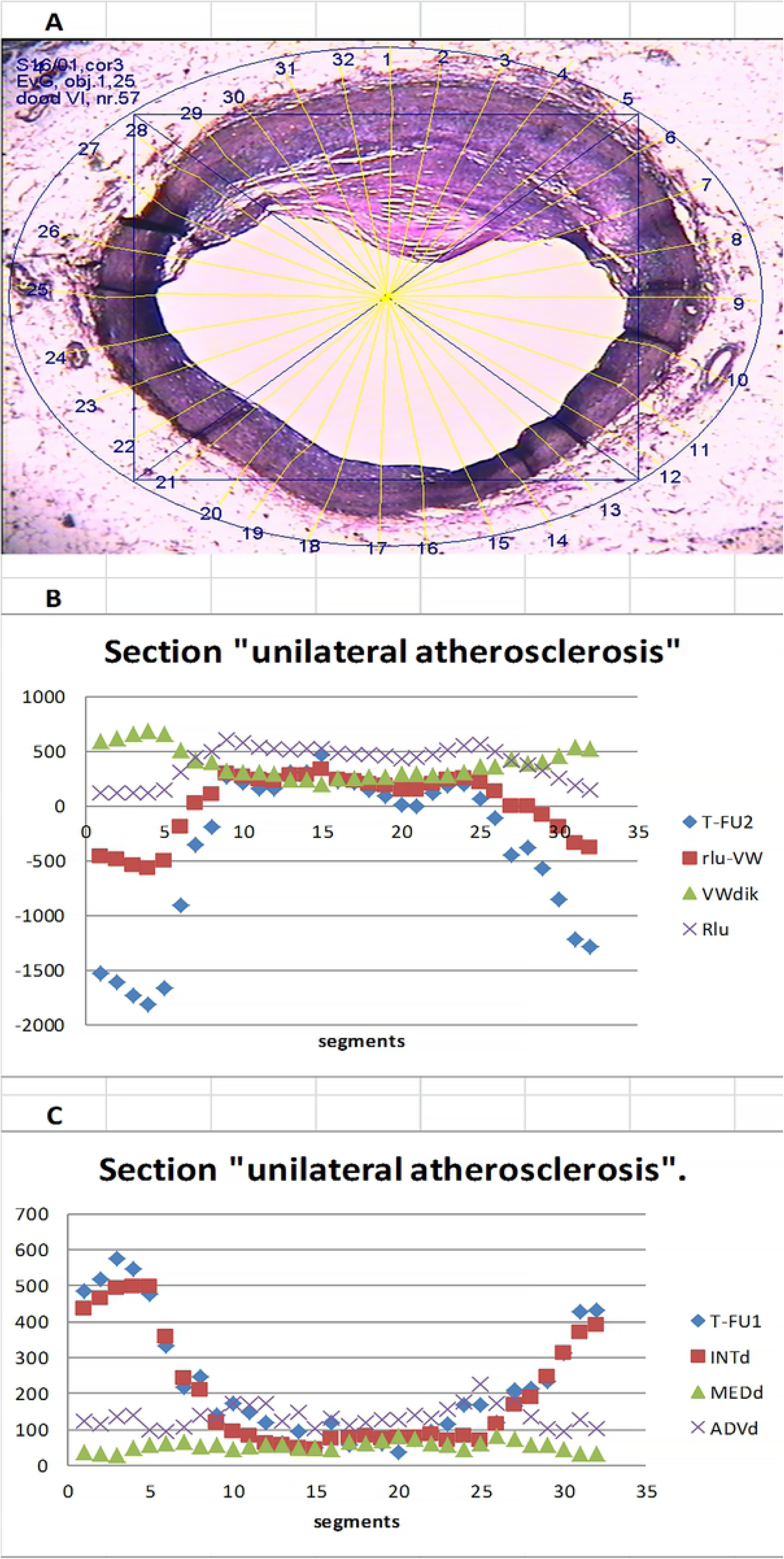
Section rated ‘unilateral atherosclerosis’. (A) Micrograph. (B) Scatter diagram T-FU2, Rlu-VWd, VWd, and Rlu vs. segment numbers. (C) T-FU1, INTd, MEDd, and ADVd vs. segment numbers.

**Fig 5.**
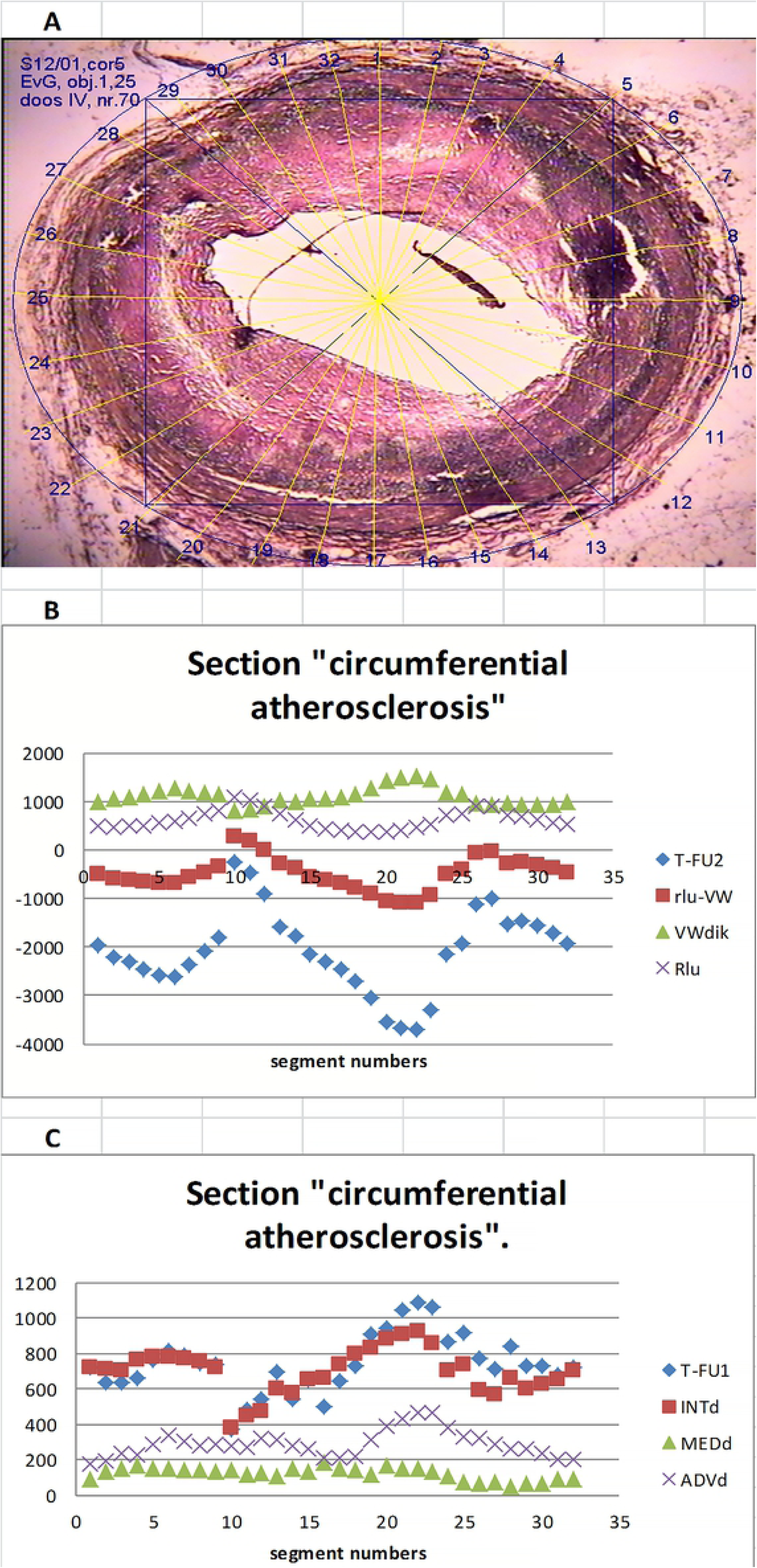
Section rated ‘circumferential atherosclerosis’. (A) Micrograph. (B) Scatter diagram T-FU2, Rlu-VWd, VWd, and Rlu vs. segment numbers. (C) T-FU1, INTd, MEDd, and ADVd vs. segment numbers.

### Triplets

Each section consisted of 16 triplets, and each triplet consisted of 20 associated data points. After data conversion to linear courses (Δ), various triplets showed that ΔT-FU2 and ΔRlu-VWd (R^2^ always > 0.95) alternated from increasing to decreasing order. Regardless of a section’s histological rating, nine triplets showed a decrease and seven triplets an increase. Increasing and decreasing triplets were not always successive.

The paths followed by ΔRlu-VWd, ΔVWd, ΔRlu, ΔINTd, ΔMEDd, and ΔADVd within each triplet were unique and differed in various triplets. Marks were discerned within several triplets (asynchronous, course-dependent, section-independent): ΔRlu-VWd ≈ 0 resulted in ΔVWd ≈ ΔRlu, ΔVWd ≈ 0 gives ΔRlu-VWd ≈ ΔRlu, and ΔRlu ≈ 0 resulted in ΔVWd ≈ −ΔRlu-VWd. Asynchronous ΔINTd, ΔMEDd, or ΔADVd ≈ 0 was also encountered without noticeable influence on the ΔVWd sequence. Even two of these being ≈ 0 did not influence the ΔVWd course. Marks in various triplets were not accompanied by ΔT-FU2 or ΔRlu-VWd similarity.

### Triplet interaction

The ΔVWd, ΔRlu, ΔINTd, ΔMEDd, and ΔADVd values of all triplets originating from each section were set against ΔRlu-VWd separately, resulting in summed trendline a-values for ΔVWd and ΔRlu = 1 (Table 5). Consecutive triplets did not constitute a systemic sequence, but ordering in descending order based on ΔVWd trendline a-values obtained a regular sequence. Three stages were observed: stage 1 marked by trendline a-values ΔVWd and ΔRlu > 0; stage 2 marked by antagonistic a-values ΔVWd and ΔRlu; stage 3 marked by a-values ΔVWd and ΔRlu < 0. Transition stage 1 to stage 2 was marked by a-value ΔVWd = 0 and by a-value ΔRlu = 1, whereas transition from stage 2 to stage 3 presented the opposite (i.e., a-value ΔVWd = 1 and a-value ΔRlu = 0). All sections showed three stages except ‘extreme unilateral atherosclerosis’, which lacked stage 1 due to a ΔVWd trendline a-value shift from stage 1 to stage 3.

**Table 5.**
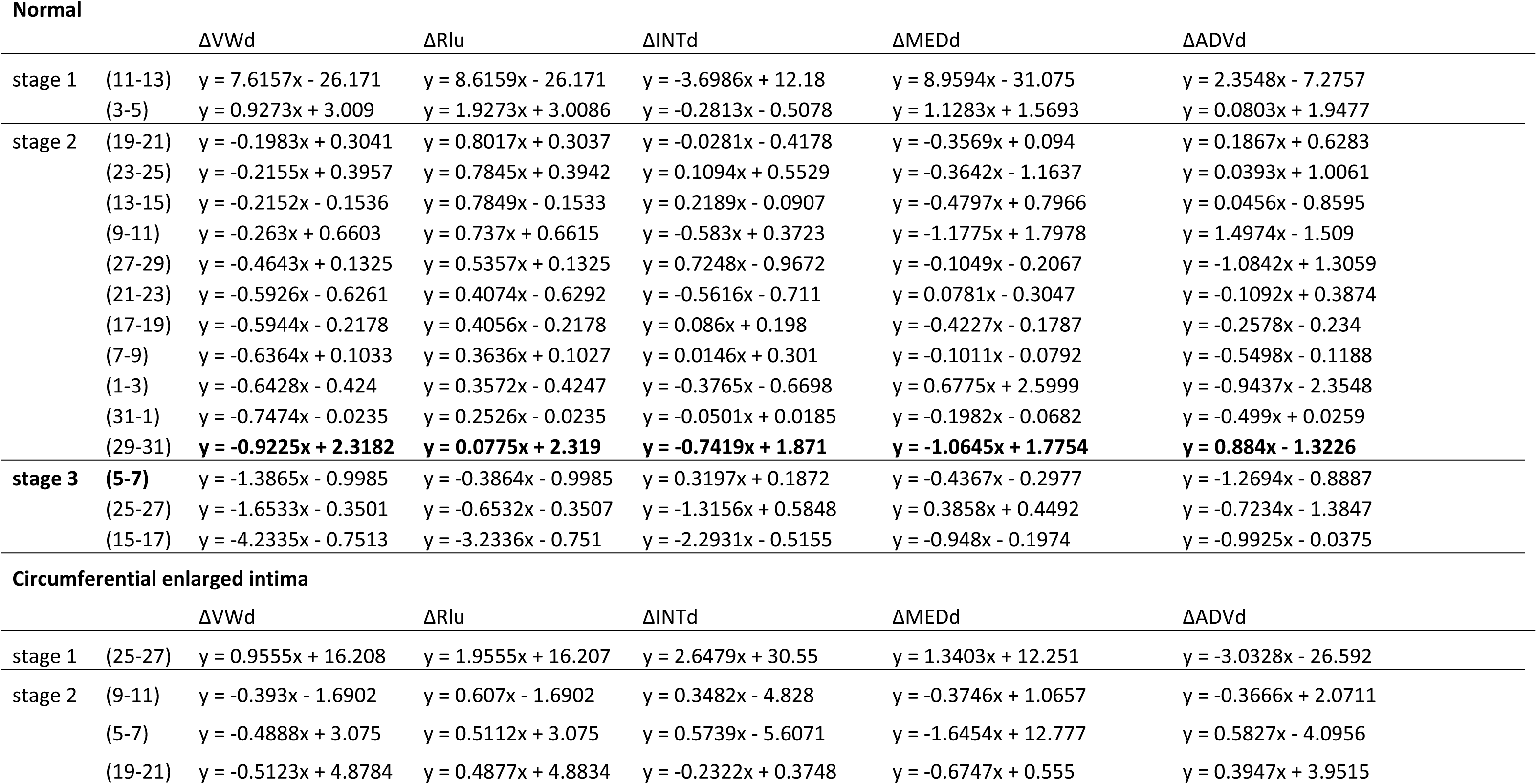
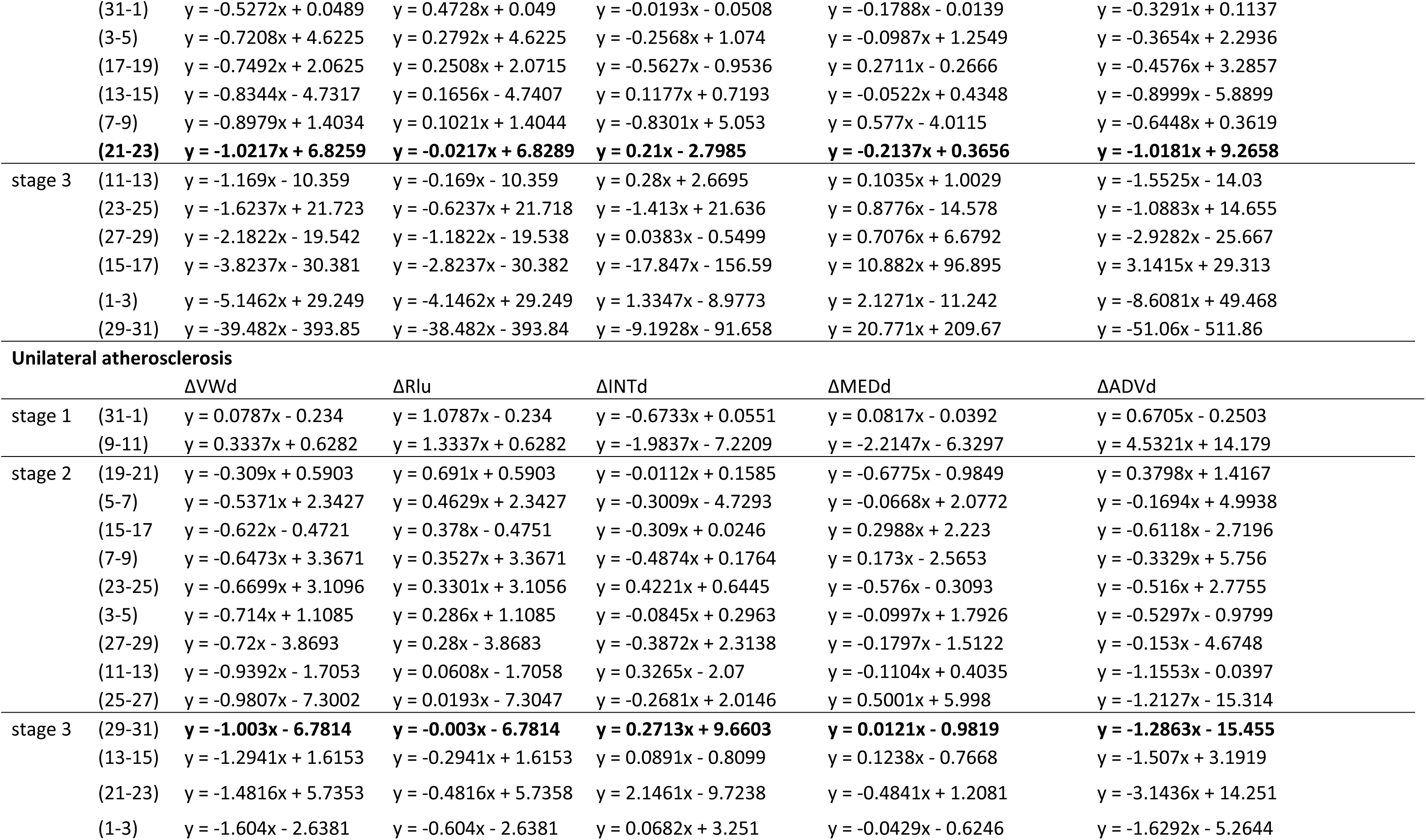
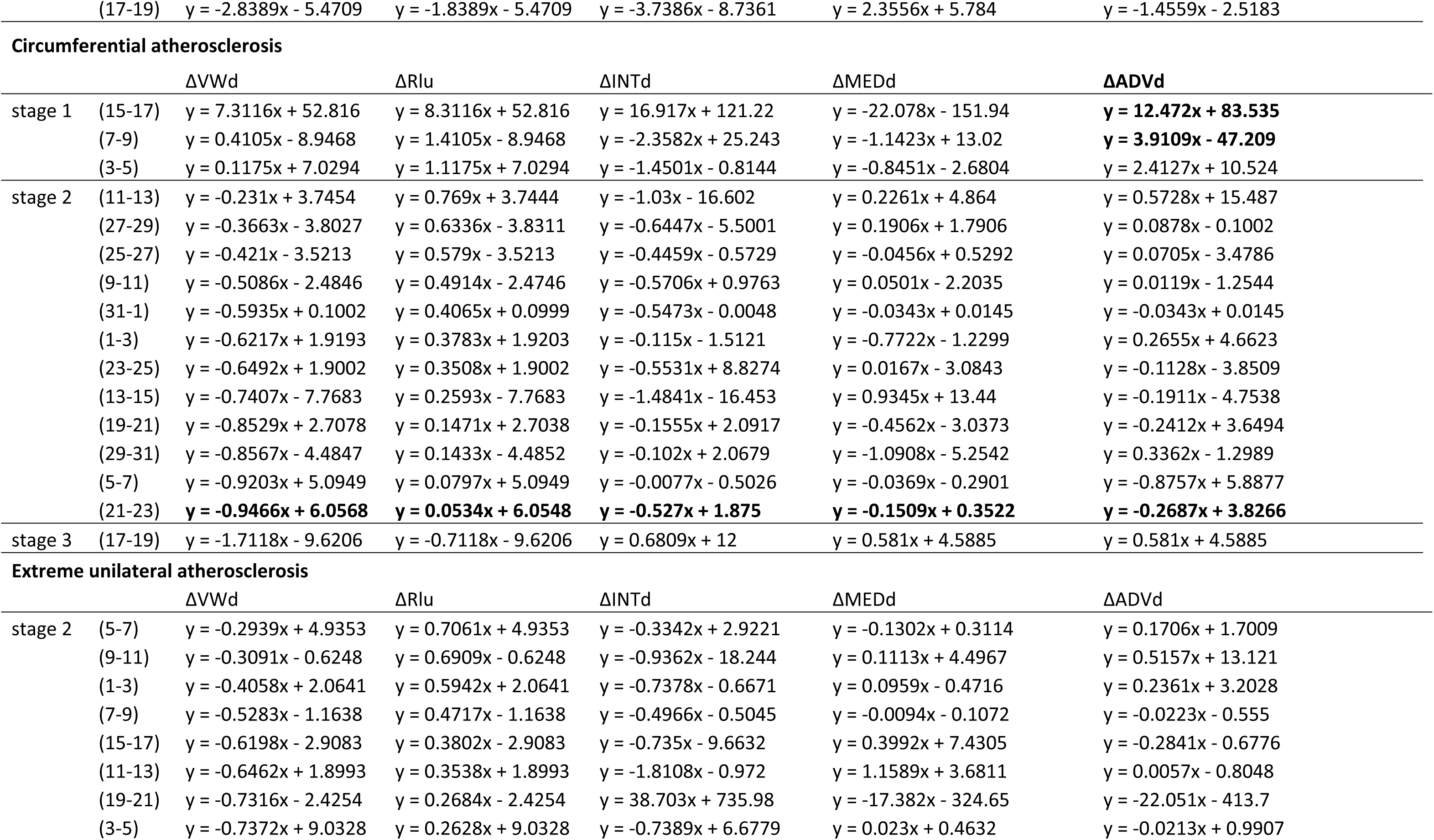
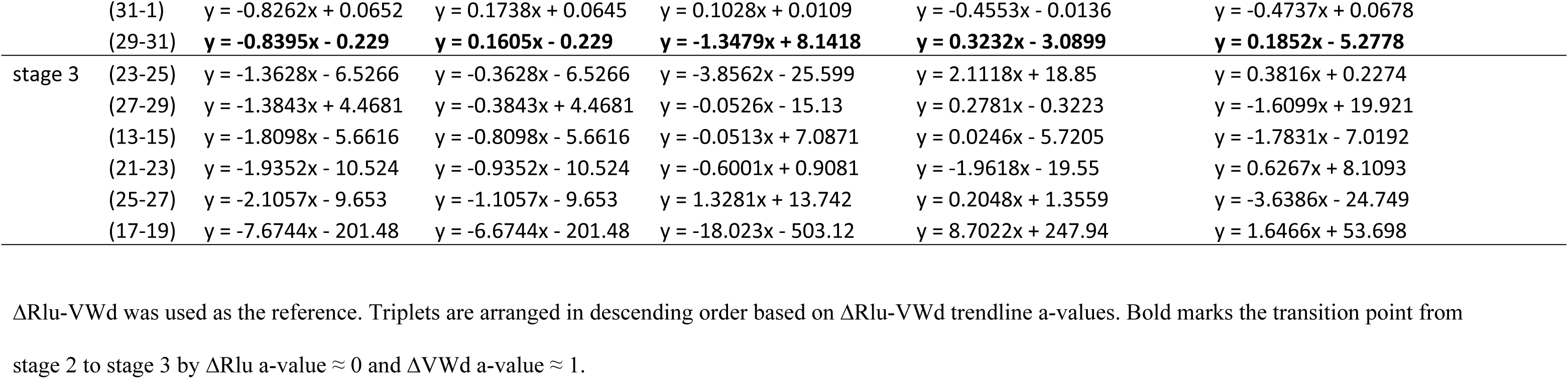
Scatter Diagram Results.

The strictly ordered course of ΔVWd and ΔRlu was not followed by a similar ΔINTd, ΔMEDd, and ΔADVd pattern; their singular courses always changed from one triplet to the next. This sometimes resulted in a parallel course with either ΔVWd or ΔRlu based on a-value similarity. Due to their asynchronous course, trendline a-value ≈ 0 was reached in different triplets.

The same ΔVWd trendline a-value in different sections regardless of pathology was accompanied by similar ΔRlu a-values but different b-values (i.e., the same direction (trendline slope) but on a different level).

### Triplet course expressed by semi-ranges

The paths followed in all sections and triplets expressed by semi-ranges (last value in triplet series minus first value) are presented in Table 6. The systems order encountered in Table 5 was no longer recognizable. Stages 1, 2, and 3 were still discerned: stage 1 defined by ΔVWd and ΔRlu semi-ranges > 0, stage 2 by antagonistic semi-ranges, and both < 0 in stage 3. Transition from stage 1 to stage 2 was not defined by semi-ranges; all values differed in various sections. The changeover from stage 2 to stage 3 was recognizable by semi-range value ΔRlu ≈ 0, accompanied by similar semi-ranges for ΔRlu-VWd and ΔVWd but antagonistic in sign. This feature was found in all sections though semi-range values differed greatly.

**Table 6.**
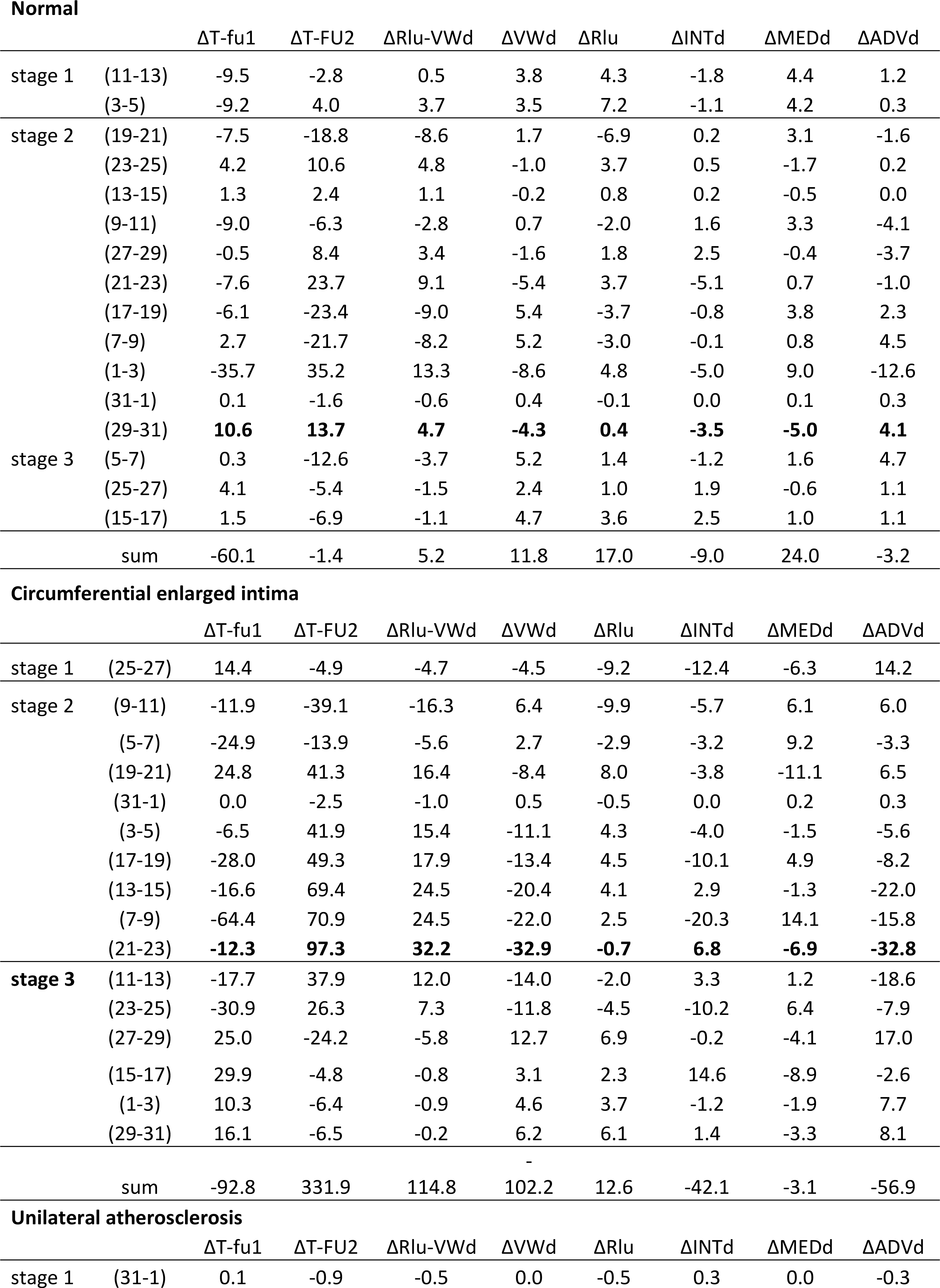
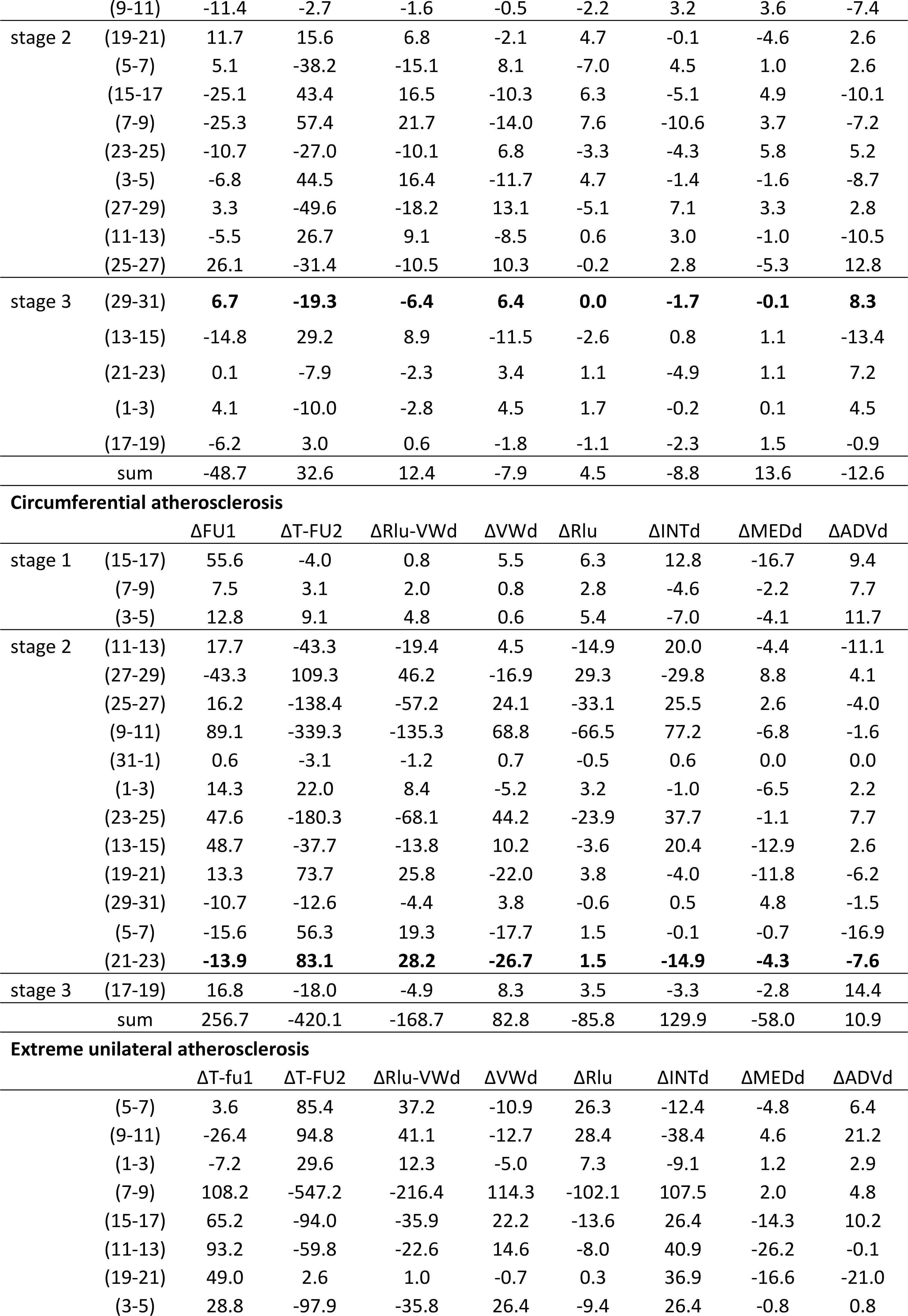
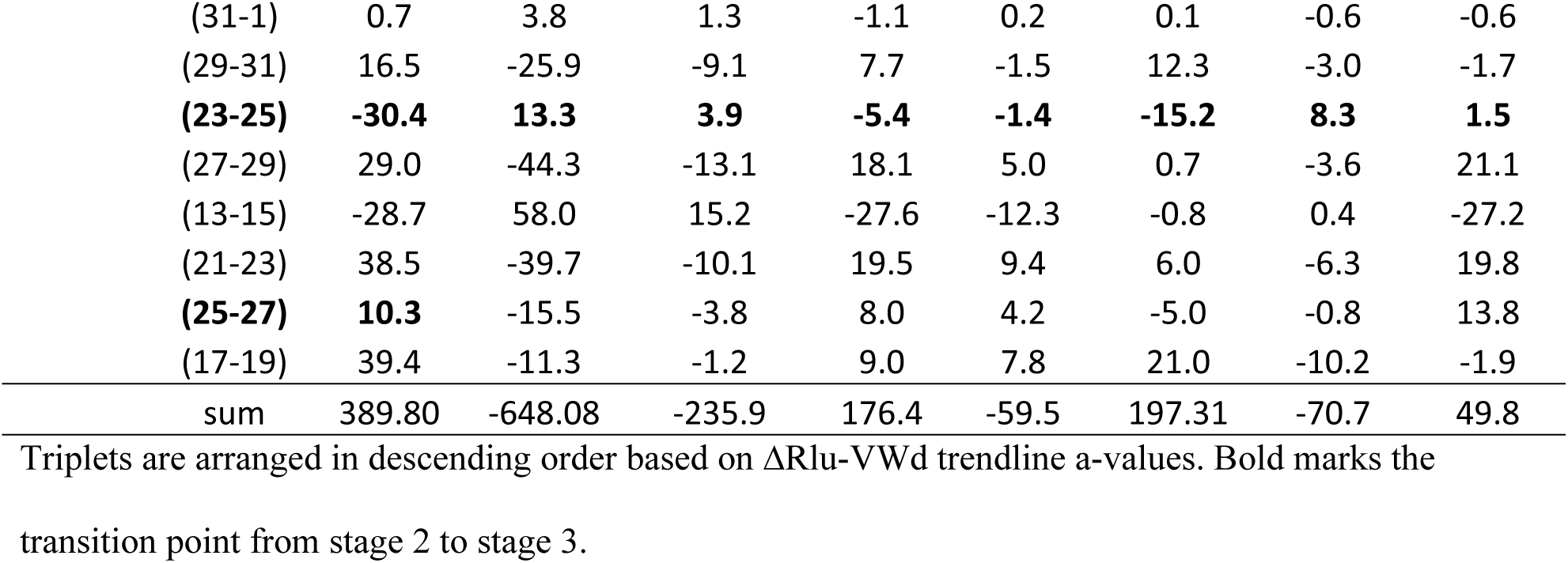
Semi-ranges Comprising the Last Value in a Triplet Series Minus the First Value.

The ΔINTd, ΔMEDd, and ΔADVd semi-ranges mutually changed from one triplet to the next, increase or decrease of one opposed to the decrease or increase of the other two, a counter-balancing system found in all sections regardless of histological rating. The summed values presented their mutual distribution: ‘normal’, semi-range ΔMEDd opposed by semi-ranges ΔINTd and ΔADVd; ‘circumferential enlarged intima’, all three had the same sign; ‘unilateral atherosclerosis’, semi-range ΔMEDd opposed by semi-ranges ΔINTd and ΔADVd; and ‘circumferential atherosclerosis’, semi-ranges ΔINTd and ΔADVd opposed by ΔMEDd (the same in ‘extreme unilateral atherosclerosis’). The last two sections had the largest semi-range ΔINTd increase together with semi-range ΔADVd growth opposed by semi-range ΔMEDd.

### Influence of ΔVW composition on ΔRlu

The individual relationship between ΔINTd, ΔMEDd, and ΔADVd and ΔRlu was not elucidated. Stage 2 differed strongly from stage 1 and stage 3, most obviously by summed values ΔVWd in ‘normal’, ‘circumferential enlarged intima’, and ‘unilateral atherosclerosis’ sections being predominantly negative due to −ΔINT, −ΔADVd, and +ΔMEDd. In contrast, ‘circumferential atherosclerosis’ sections presented predominantly positive ΔVWd values due to +ΔINTd, −ΔMEDd, and −ΔADVd, whereas ‘extreme unilateral atherosclerosis’ had positive ΔVWd values due to +ΔINTd, −ΔMEDd, and +ΔADVd.

Comparisons of semi-range ratios ΔINT, ΔMEDd, or ΔADVd to ΔVWd and ΔRlu showed variability in the single semi-range values (Table 7). Naturally, the summed semi-range values ΔINTd, ΔMEDd, and ΔADVd in proportion to semi-range ΔVWd equaled 1, but in proportion to semi-range ΔRlu a systemic course developed similar to the semi-range ratio of ΔVWd to ΔRlu. These results were found in all sections independent of semi-range ΔINTd, ΔMEDd, or ΔADVd values.

**Table 7.**
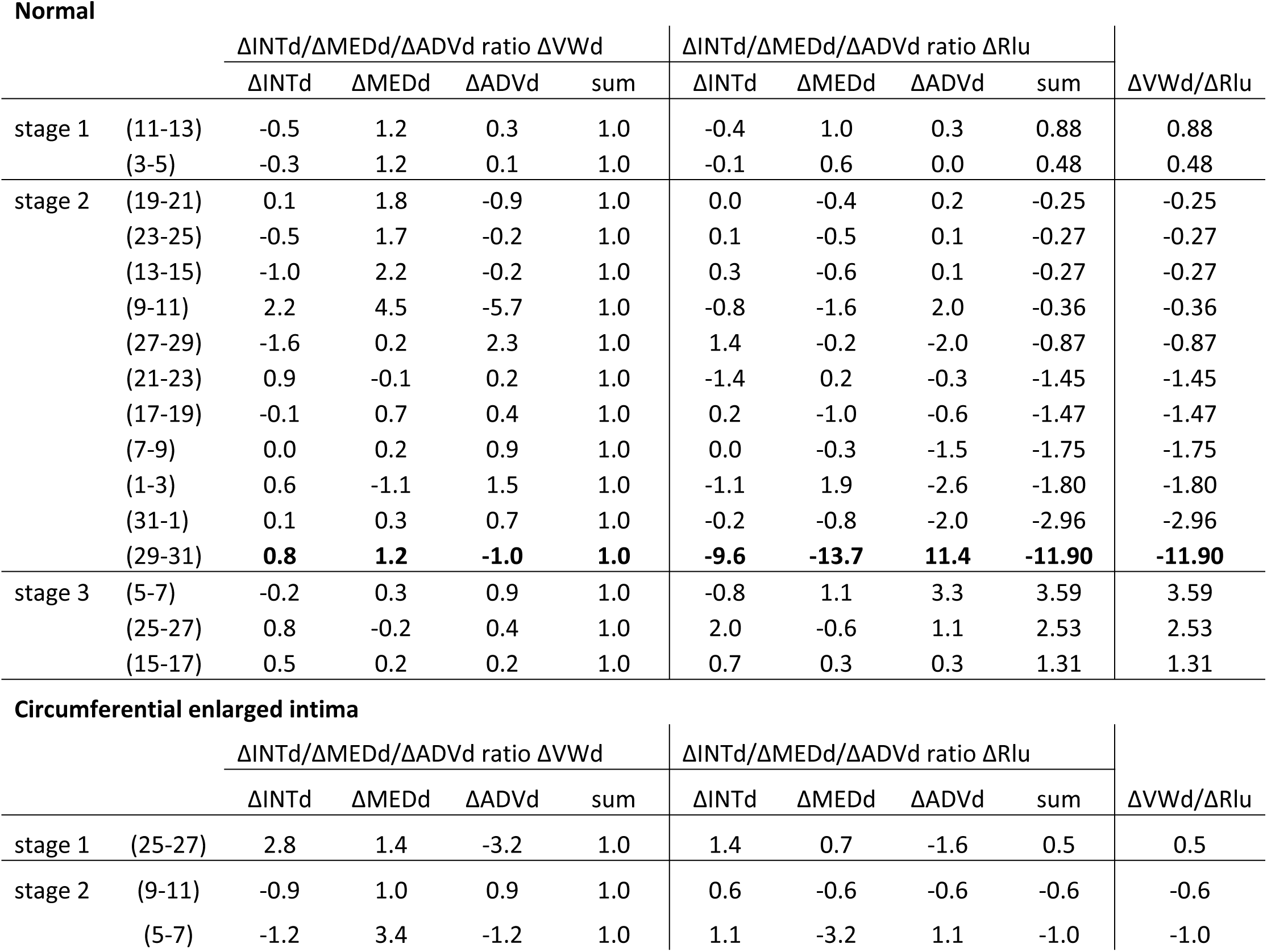
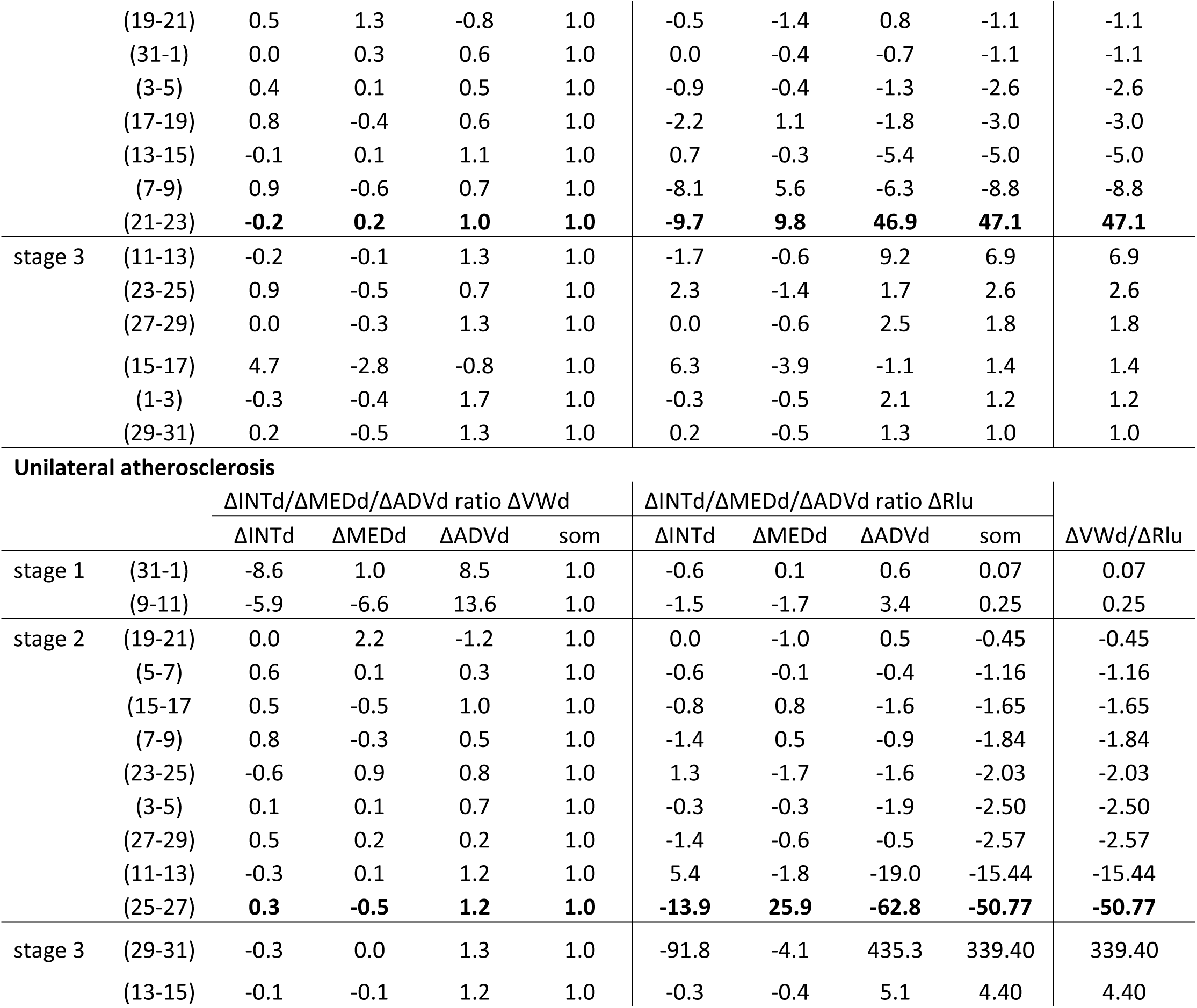
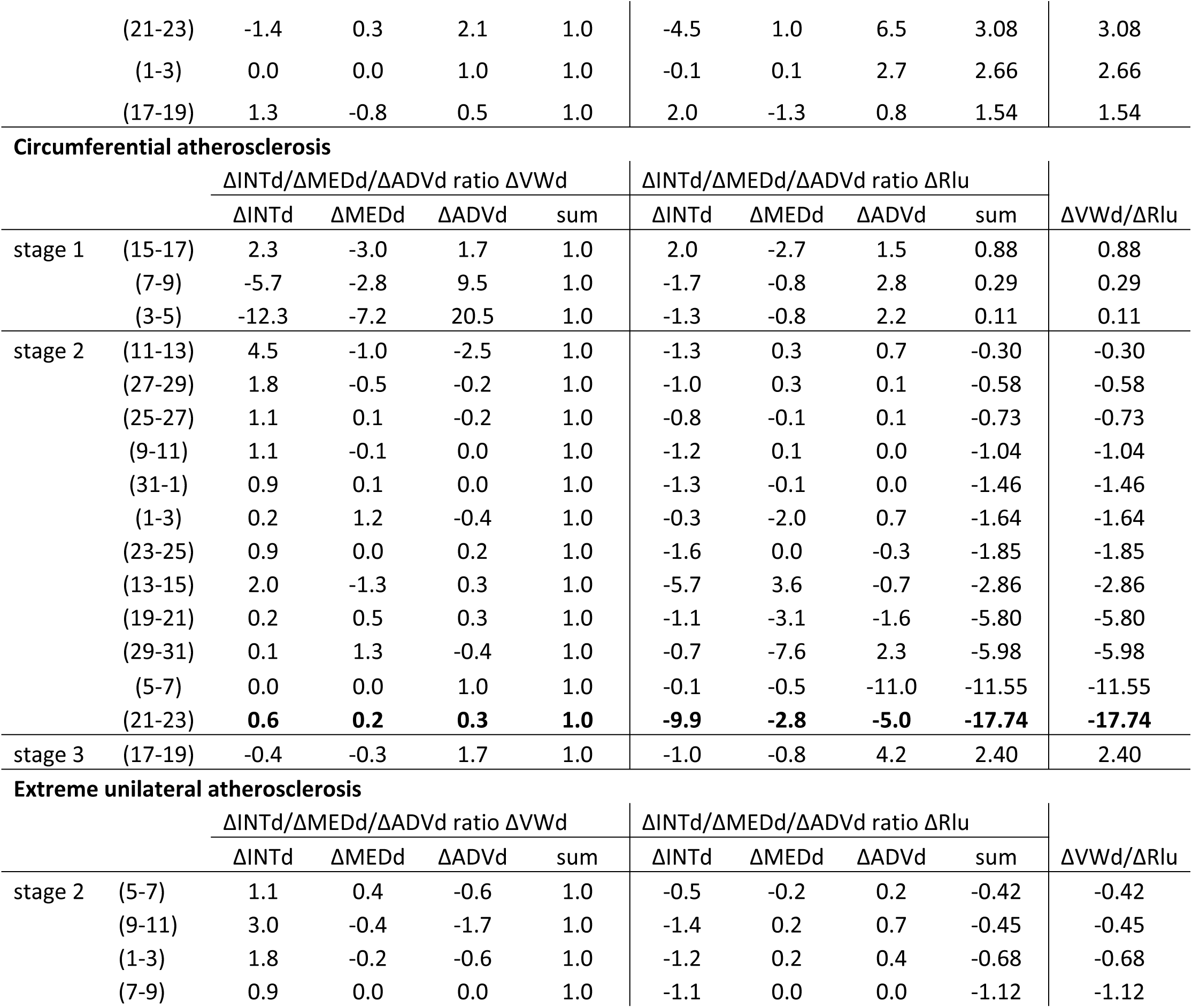
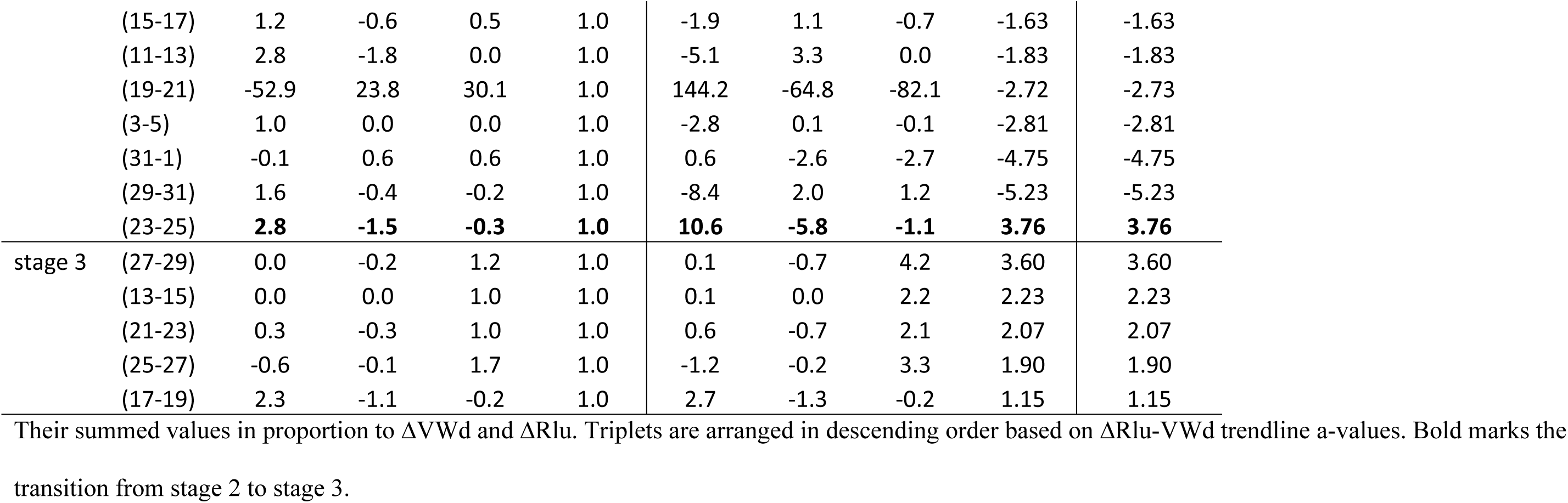
Semi-ranges ΔINTd, ΔMEDd, and ΔADVd Expressed in Proportion to ΔVWd and ΔRlu.

| | | ΔINTd/ΔMEDd/ΔADVd ratio ΔVWd | | | | ΔINTd/ΔMEDd/ΔADVd ratio ΔRlu | | | | ΔVWd/ΔRlu |
| --- | --- | --- | --- | --- | --- | --- | --- | --- | --- | --- |
| | | ΔINTd | ΔMEDd | ΔADVd | sum | ΔINTd | ΔMEDd | ΔADVd | sum | |
| stage 1 | (11-13) | -0.5 | 1.2 | 0.3 | 1.0 | -0.4 | 1.0 | 0.3 | 0.88 | 0.88 |
|  | (3-5) | -0.3 | 1.2 | 0.1 | 1.0 | -0.1 | 0.6 | 0.0 | 0.48 | 0.48 |
| stage 2 | (19-21) | 0.1 | 1.8 | -0.9 | 1.0 | 0.0 | -0.4 | 0.2 | -0.25 | -0.25 |
|  | (23-25) | -0.5 | 1.7 | -0.2 | 1.0 | 0.1 | -0.5 | 0.1 | -0.27 | -0.27 |
|  | (13-15) | -1.0 | 2.2 | -0.2 | 1.0 | 0.3 | -0.6 | 0.1 | -0.27 | -0.27 |
|  | (9-11) | 2.2 | 4.5 | -5.7 | 1.0 | -0.8 | -1.6 | 2.0 | -0.36 | -0.36 |
|  | (27-29) | -1.6 | 0.2 | 2.3 | 1.0 | 1.4 | -0.2 | -2.0 | -0.87 | -0.87 |
|  | (21-23) | 0.9 | -0.1 | 0.2 | 1.0 | -1.4 | 0.2 | -0.3 | -1.45 | -1.45 |
|  | (17-19) | -0.1 | 0.7 | 0.4 | 1.0 | 0.2 | -1.0 | -0.6 | -1.47 | -1.47 |
|  | (7-9) | 0.0 | 0.2 | 0.9 | 1.0 | 0.0 | -0.3 | -1.5 | -1.75 | -1.75 |
|  | (1-3) | 0.6 | -1.1 | 1.5 | 1.0 | -1.1 | 1.9 | -2.6 | -1.80 | -1.80 |
|  | (31-1) | 0.1 | 0.3 | 0.7 | 1.0 | -0.2 | -0.8 | -2.0 | -2.96 | -2.96 |
|  | (29-31) | **0.8** | **1.2** | **-1.0** | **1.0** | **-9.6** | **-13.7** | **11.4** | **-11.90** | **-11.90** |
| stage 3 | (5-7) | -0.2 | 0.3 | 0.9 | 1.0 | -0.8 | 1.1 | 3.3 | 3.59 | 3.59 |
|  | (25-27) | 0.8 | -0.2 | 0.4 | 1.0 | 2.0 | -0.6 | 1.1 | 2.53 | 2.53 |
|  | (15-17) | 0.5 | 0.2 | 0.2 | 1.0 | 0.7 | 0.3 | 0.3 | 1.31 | 1.31 |

**Circumferential enlarged intima**
| | | ΔINTd/ΔMEDd/ΔADVd ratio ΔVWd | | | | ΔINTd/ΔMEDd/ΔADVd ratio ΔRlu | | | | ΔVWd/ΔRlu |
| --- | --- | --- | --- | --- | --- | --- | --- | --- | --- | --- |
| | | ΔINTd | ΔMEDd | ΔADVd | sum | ΔINTd | ΔMEDd | ΔADVd | sum | |
| stage 1 | (25-27) | 2.8 | 1.4 | -3.2 | 1.0 | 1.4 | 0.7 | -1.6 | 0.5 | 0.5 |
| stage 2 | (9-11) | -0.9 | 1.0 | 0.9 | 1.0 | 0.6 | -0.6 | -0.6 | -0.6 | -0.6 |
|  | (5-7) | -1.2 | 3.4 | -1.2 | 1.0 | 1.1 | -3.2 | 1.1 | -1.0 | -1.0 |
|  | (19-21) | 0.5 | 1.3 | -0.8 | 1.0 | -0.5 | -1.4 | 0.8 | -1.1 | -1.1 |
|  | (31-1) | 0.0 | 0.3 | 0.6 | 1.0 | 0.0 | -0.4 | -0.7 | -1.1 | -1.1 |
|  | (3-5) | 0.4 | 0.1 | 0.5 | 1.0 | -0.9 | -0.4 | -1.3 | -2.6 | -2.6 |
|  | (17-19) | 0.8 | -0.4 | 0.6 | 1.0 | -2.2 | 1.1 | -1.8 | -3.0 | -3.0 |
|  | (13-15) | -0.1 | 0.1 | 1.1 | 1.0 | 0.7 | -0.3 | -5.4 | -5.0 | -5.0 |
|  | (7-9) | 0.9 | -0.6 | 0.7 | 1.0 | -8.1 | 5.6 | -6.3 | -8.8 | -8.8 |
|  | (21-23) | **-0.2** | **0.2** | **1.0** | **1.0** | **-9.7** | **9.8** | **46.9** | **47.1** | **47.1** |
| stage 3 | (11-13) | -0.2 | -0.1 | 1.3 | 1.0 | -1.7 | -0.6 | 9.2 | 6.9 | 6.9 |
|  | (23-25) | 0.9 | -0.5 | 0.7 | 1.0 | 2.3 | -1.4 | 1.7 | 2.6 | 2.6 |
|  | (27-29) | 0.0 | -0.3 | 1.3 | 1.0 | 0.0 | -0.6 | 2.5 | 1.8 | 1.8 |
|  | (15-17) | 4.7 | -2.8 | -0.8 | 1.0 | 6.3 | -3.9 | -1.1 | 1.4 | 1.4 |
|  | (1-3) | -0.3 | -0.4 | 1.7 | 1.0 | -0.3 | -0.5 | 2.1 | 1.2 | 1.2 |
|  | (29-31) | 0.2 | -0.5 | 1.3 | 1.0 | 0.2 | -0.5 | 1.3 | 1.0 | 1.0 |

Unilateral atherosclerosis
| | | $\Delta\text{INTd}/\Delta\text{MEDd}/\Delta\text{ADVd}$ ratio $\Delta\text{VWd}$ | | | | $\Delta\text{INTd}/\Delta\text{MEDd}/\Delta\text{ADVd}$ ratio $\Delta\text{Rlu}$ | | | | $\Delta\text{VWd}/\Delta\text{Rlu}$ |
| --- | --- | --- | --- | --- | --- | --- | --- | --- | --- | --- |
| | | $\Delta\text{INTd}$ | $\Delta\text{MEDd}$ | $\Delta\text{ADVd}$ | som | $\Delta\text{INTd}$ | $\Delta\text{MEDd}$ | $\Delta\text{ADVd}$ | som | |
| stage 1 | (31-1) | -8.6 | 1.0 | 8.5 | 1.0 | -0.6 | 0.1 | 0.6 | 0.07 | 0.07 |
|  | (9-11) | -5.9 | -6.6 | 13.6 | 1.0 | -1.5 | -1.7 | 3.4 | 0.25 | 0.25 |
| stage 2 | (19-21) | 0.0 | 2.2 | -1.2 | 1.0 | 0.0 | -1.0 | 0.5 | -0.45 | -0.45 |
|  | (5-7) | 0.6 | 0.1 | 0.3 | 1.0 | -0.6 | -0.1 | -0.4 | -1.16 | -1.16 |
|  | (15-17) | 0.5 | -0.5 | 1.0 | 1.0 | -0.8 | 0.8 | -1.6 | -1.65 | -1.65 |
|  | (7-9) | 0.8 | -0.3 | 0.5 | 1.0 | -1.4 | 0.5 | -0.9 | -1.84 | -1.84 |
|  | (23-25) | -0.6 | 0.9 | 0.8 | 1.0 | 1.3 | -1.7 | -1.6 | -2.03 | -2.03 |
|  | (3-5) | 0.1 | 0.1 | 0.7 | 1.0 | -0.3 | -0.3 | -1.9 | -2.50 | -2.50 |
|  | (27-29) | 0.5 | 0.2 | 0.2 | 1.0 | -1.4 | -0.6 | -0.5 | -2.57 | -2.57 |
|  | (11-13) | -0.3 | 0.1 | 1.2 | 1.0 | 5.4 | -1.8 | -19.0 | -15.44 | -15.44 |
|  | (25-27) | **0.3** | **-0.5** | **1.2** | **1.0** | **-13.9** | **25.9** | **-62.8** | **-50.77** | **-50.77** |
| stage 3 | (29-31) | -0.3 | 0.0 | 1.3 | 1.0 | -91.8 | -4.1 | 435.3 | 339.40 | 339.40 |
|  | (13-15) | -0.1 | -0.1 | 1.2 | 1.0 | -0.3 | -0.4 | 5.1 | 4.40 | 4.40 |

|  |  |  |  |  |  |  |  |  |  |
| --- | --- | --- | --- | --- | --- | --- | --- | --- | --- |
| (21-23) | -1.4 | 0.3 | 2.1 | 1.0 | -4.5 | 1.0 | 6.5 | 3.08 | 3.08 |
| (1-3) | 0.0 | 0.0 | 1.0 | 1.0 | -0.1 | 0.1 | 2.7 | 2.66 | 2.66 |
| (17-19) | 1.3 | -0.8 | 0.5 | 1.0 | 2.0 | -1.3 | 0.8 | 1.54 | 1.54 |

Circumferential atherosclerosis
| | | $\Delta\text{INTd}/\Delta\text{MEDd}/\Delta\text{ADVd}$ ratio $\Delta\text{VWd}$ | | | | $\Delta\text{INTd}/\Delta\text{MEDd}/\Delta\text{ADVd}$ ratio $\Delta\text{Rlu}$ | | | | $\Delta\text{VWd}/\Delta\text{Rlu}$ |
| --- | --- | --- | --- | --- | --- | --- | --- | --- | --- | --- |
| | | $\Delta\text{INTd}$ | $\Delta\text{MEDd}$ | $\Delta\text{ADVd}$ | sum | $\Delta\text{INTd}$ | $\Delta\text{MEDd}$ | $\Delta\text{ADVd}$ | sum | |
| stage 1 | (15-17) | 2.3 | -3.0 | 1.7 | 1.0 | 2.0 | -2.7 | 1.5 | 0.88 | 0.88 |
|  | (7-9) | -5.7 | -2.8 | 9.5 | 1.0 | -1.7 | -0.8 | 2.8 | 0.29 | 0.29 |
|  | (3-5) | -12.3 | -7.2 | 20.5 | 1.0 | -1.3 | -0.8 | 2.2 | 0.11 | 0.11 |
| stage 2 | (11-13) | 4.5 | -1.0 | -2.5 | 1.0 | -1.3 | 0.3 | 0.7 | -0.30 | -0.30 |
|  | (27-29) | 1.8 | -0.5 | -0.2 | 1.0 | -1.0 | 0.3 | 0.1 | -0.58 | -0.58 |
|  | (25-27) | 1.1 | 0.1 | -0.2 | 1.0 | -0.8 | -0.1 | 0.1 | -0.73 | -0.73 |
|  | (9-11) | 1.1 | -0.1 | 0.0 | 1.0 | -1.2 | 0.1 | 0.0 | -1.04 | -1.04 |
|  | (31-1) | 0.9 | 0.1 | 0.0 | 1.0 | -1.3 | -0.1 | 0.0 | -1.46 | -1.46 |
|  | (1-3) | 0.2 | 1.2 | -0.4 | 1.0 | -0.3 | -2.0 | 0.7 | -1.64 | -1.64 |
|  | (23-25) | 0.9 | 0.0 | 0.2 | 1.0 | -1.6 | 0.0 | -0.3 | -1.85 | -1.85 |
|  | (13-15) | 2.0 | -1.3 | 0.3 | 1.0 | -5.7 | 3.6 | -0.7 | -2.86 | -2.86 |
|  | (19-21) | 0.2 | 0.5 | 0.3 | 1.0 | -1.1 | -3.1 | -1.6 | -5.80 | -5.80 |
|  | (29-31) | 0.1 | 1.3 | -0.4 | 1.0 | -0.7 | -7.6 | 2.3 | -5.98 | -5.98 |
|  | (5-7) | 0.0 | 0.0 | 1.0 | 1.0 | -0.1 | -0.5 | -11.0 | -11.55 | -11.55 |
|  | (21-23) | **0.6** | **0.2** | **0.3** | **1.0** | **-9.9** | **-2.8** | **-5.0** | **-17.74** | **-17.74** |
| stage 3 | (17-19) | -0.4 | -0.3 | 1.7 | 1.0 | -1.0 | -0.8 | 4.2 | 2.40 | 2.40 |

Extreme unilateral atherosclerosis
| | | $\Delta\text{INTd}/\Delta\text{MEDd}/\Delta\text{ADVd}$ ratio $\Delta\text{VWd}$ | | | | $\Delta\text{INTd}/\Delta\text{MEDd}/\Delta\text{ADVd}$ ratio $\Delta\text{Rlu}$ | | | | $\Delta\text{VWd}/\Delta\text{Rlu}$ |
| --- | --- | --- | --- | --- | --- | --- | --- | --- | --- | --- |
| | | $\Delta\text{INTd}$ | $\Delta\text{MEDd}$ | $\Delta\text{ADVd}$ | sum | $\Delta\text{INTd}$ | $\Delta\text{MEDd}$ | $\Delta\text{ADVd}$ | sum | |
| stage 2 | (5-7) | 1.1 | 0.4 | -0.6 | 1.0 | -0.5 | -0.2 | 0.2 | -0.42 | -0.42 |
|  | (9-11) | 3.0 | -0.4 | -1.7 | 1.0 | -1.4 | 0.2 | 0.7 | -0.45 | -0.45 |
|  | (1-3) | 1.8 | -0.2 | -0.6 | 1.0 | -1.2 | 0.2 | 0.4 | -0.68 | -0.68 |
|  | (7-9) | 0.9 | 0.0 | 0.0 | 1.0 | -1.1 | 0.0 | 0.0 | -1.12 | -1.12 |
|  | (15-17) | 1.2 | -0.6 | 0.5 | 1.0 | -1.9 | 1.1 | -0.7 | -1.63 | -1.63 |
|  | (11-13) | 2.8 | -1.8 | 0.0 | 1.0 | -5.1 | 3.3 | 0.0 | -1.83 | -1.83 |
|  | (19-21) | -52.9 | 23.8 | 30.1 | 1.0 | 144.2 | -64.8 | -82.1 | -2.72 | -2.73 |
|  | (3-5) | 1.0 | 0.0 | 0.0 | 1.0 | -2.8 | 0.1 | -0.1 | -2.81 | -2.81 |
|  | (31-1) | -0.1 | 0.6 | 0.6 | 1.0 | 0.6 | -2.6 | -2.7 | -4.75 | -4.75 |
|  | (29-31) | 1.6 | -0.4 | -0.2 | 1.0 | -8.4 | 2.0 | 1.2 | -5.23 | -5.23 |
|  | (23-25) | **2.8** | **-1.5** | **-0.3** | **1.0** | **10.6** | **-5.8** | **-1.1** | **3.76** | **3.76** |
| stage 3 | (27-29) | 0.0 | -0.2 | 1.2 | 1.0 | 0.1 | -0.7 | 4.2 | 3.60 | 3.60 |
|  | (13-15) | 0.0 | 0.0 | 1.0 | 1.0 | 0.1 | 0.0 | 2.2 | 2.23 | 2.23 |
|  | (21-23) | 0.3 | -0.3 | 1.0 | 1.0 | 0.6 | -0.7 | 2.1 | 2.07 | 2.07 |
|  | (25-27) | -0.6 | -0.1 | 1.7 | 1.0 | -1.2 | -0.2 | 3.3 | 1.90 | 1.90 |
|  | (17-19) | 2.3 | -1.1 | -0.2 | 1.0 | 2.7 | -1.3 | -0.2 | 1.15 | 1.15 |

Their summed values in proportion to $\Delta V W d$ and $\Delta R l u$ . Triplets are arranged in descending order based on $\Delta R l u$ - $V W d$ trendline a-values. Bold marks the
transition from stage 2 to stage 3.

Despite semi-range ΔINTd, ΔMEDd, and ΔADVd mutually variable interactions, semi-range ΔVWd was strongly associated with semi-range ΔRlu in a systemic way (Table 8). This finding emphasizes that semi-ranges reliably reproduce measurement data.

**Table 8.**
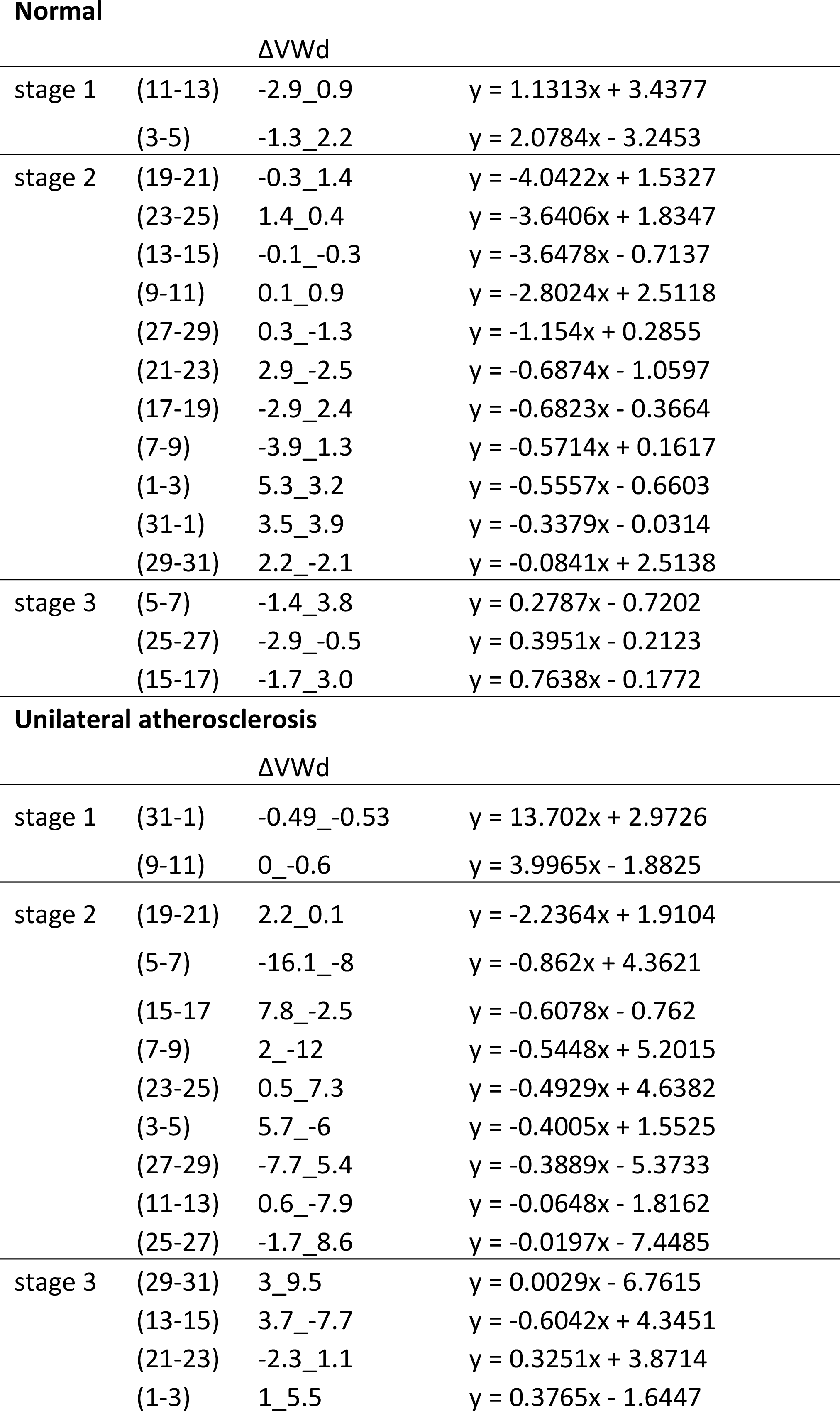
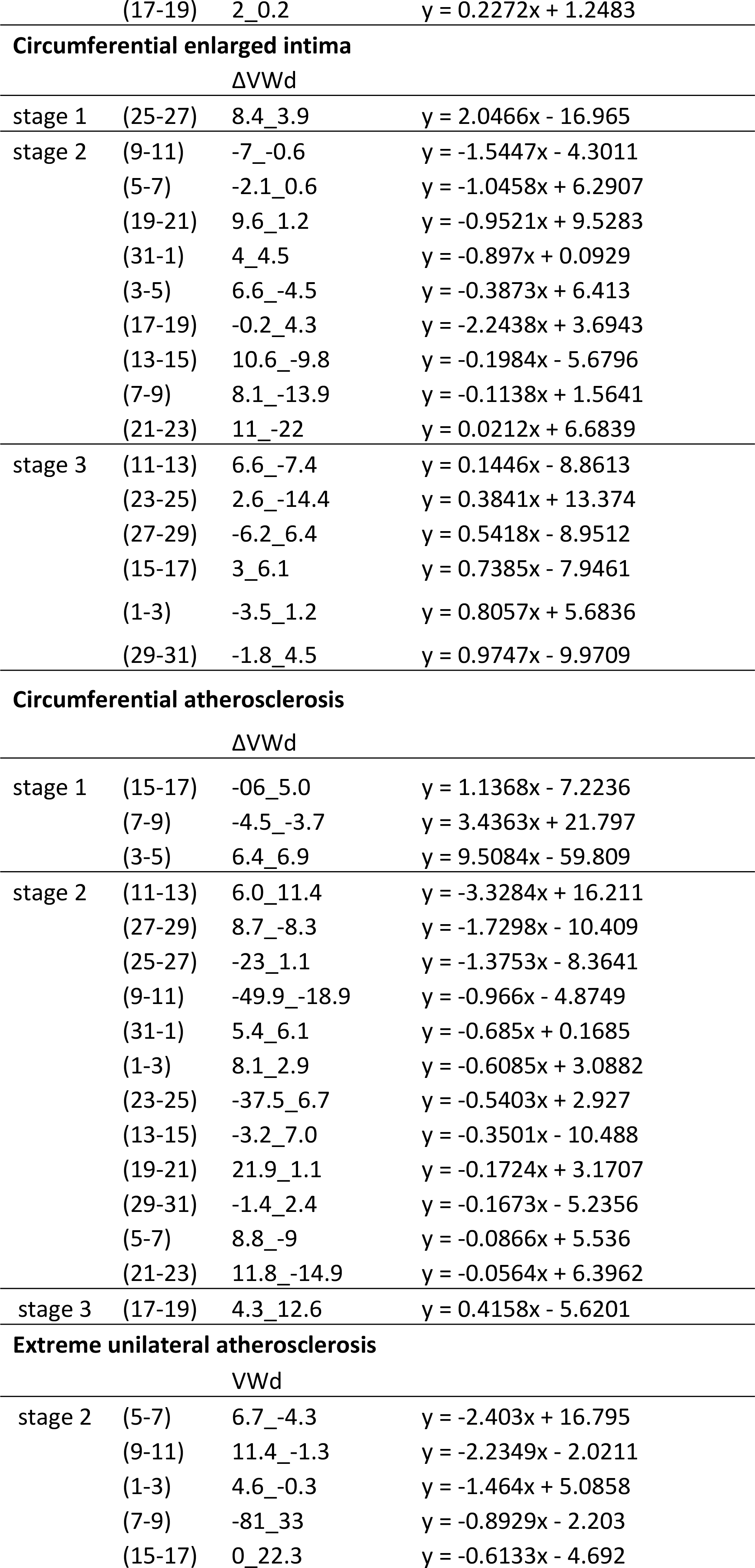
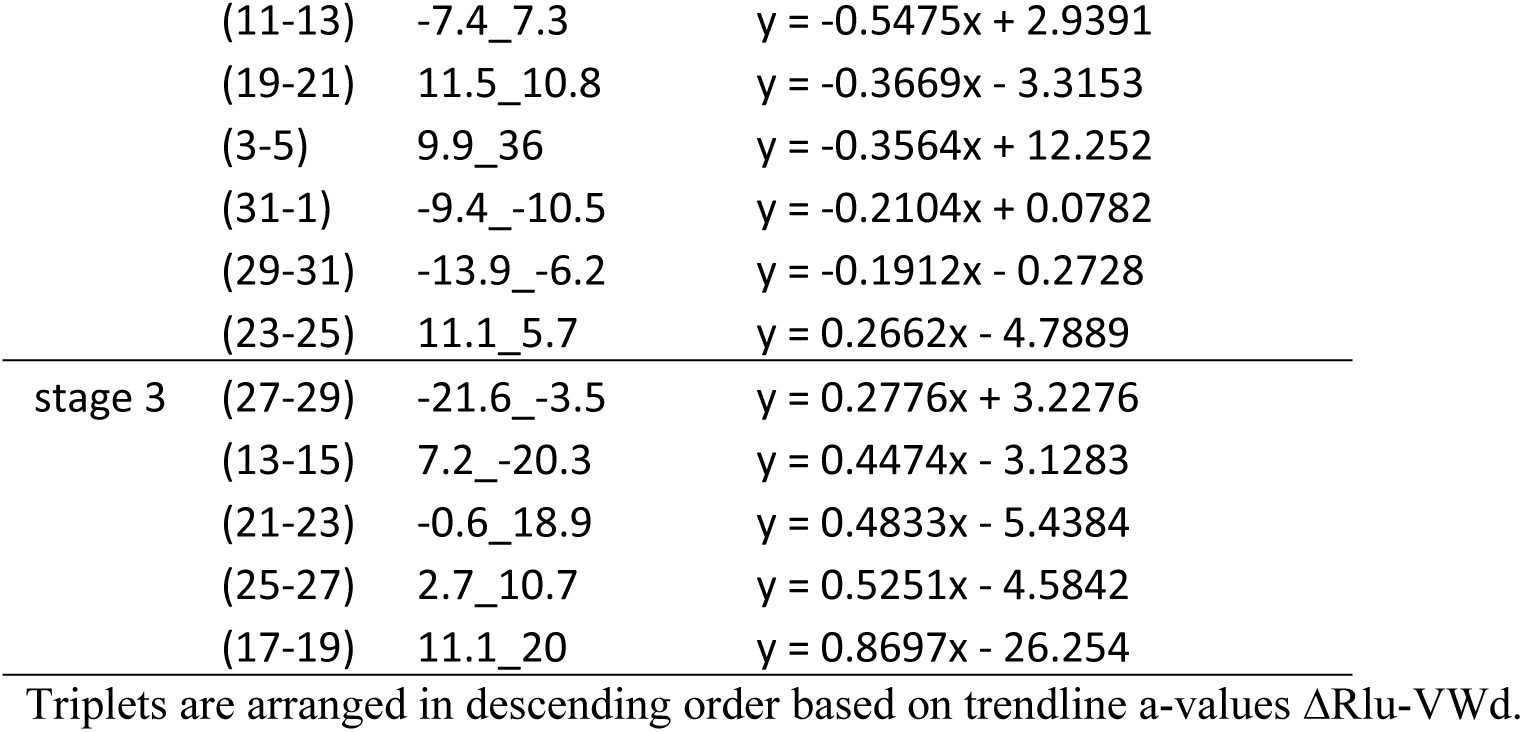
Trendline Formula ΔRlu vs. ΔVWd Triplets.

## Discussion

Excision of pressure perfused vessels is a standard method used to prepare and examine the cardiac vessels at autopsy because it allows better approximation of the *invivo* lumen and vessel wall configuration. However, no agreement has been reached on the best post-mortem vessel examination technique, with wide ranges of perfusion pressures used in different studies making it impossible to compare results. Here, we used non-perfused vessels to better preserve the elastic characteristic of the coronary vessel wall, with residual wall stress clearly evident by the cut ends of the walls moving away from each other after longitudinal incision. Sections cut perpendicular to the longitudinal axis maintain residual wall stress, thereby retaining the basal lumen and vessel wall configuration [7]. This approach preserves the mutual relationships, as confirmed by our results.

In essence, the main basic process is the ‘counter-balancing’ system: growth of ΔINTd, ΔMEDd, or ΔADVd counter-balanced by the other two. As such, these are functionally united in a direct way (ΔFU-1). This basic function stays intact regardless of vessel wall pathology, though triplets present the same sign in a minority of sections. In an indirect way ΔINTd, ΔMEDd, and ΔADVd determine the course of ΔVWd in every triplet (increasing or decreasing), its measure by semi-range values and their sign change.

Within each section, stages are defined in which the mutual relationship between ΔVWd and ΔRlu changes, from an increase in ΔVWd and increased ΔRlu (stage 1 and stage 3) to a decrease in ΔRlu and increased ΔVWd (stage 2). The transition of stage 1 to stage 2 is marked by trendline a-value ΔVWd = 0 and ΔRlu = 1, whereas the transition from stage 2 to stage 3 is marked by a-value ΔVWd =1 and ΔRlu=0. The a-value ΔVWd = 0 and ΔRlu=0 reflect the peak/bottom parabola values before data transition to Δ values, which transforms a parabolic course to a linear course. As such, stages are defined by the ΔVWd and ΔRlu parabolic course remnants, a change from synchronic (same trendline a-value sign, stages 1 and 3) to asynchronic (opposite trendline sign, stage 2). This proves that the ΔVWd/ΔRlu relationship is course- and sign-dependent and, therefore, independent of their values in absolute terms.

Atherosclerotic and non-atherosclerotic sections cannot be differentiated by triplet behavior. For example, ΔINTd growth with atherosclerosis is marked only by a shift from stage 1 to stage 3. This illustrates that even atherosclerotic sections exhibit an increase in ΔRlu in relation to ΔVWd growth, which implies independence from vessel wall constitution. Support for this view is given by the summed semi-range values ΔINTd, ΔMEDd, and ΔADVd in proportion to semi-range ΔRlu, which is similar to the ΔVWd/ΔRlu ratio in absolute terms. Therefore, the atherosclerotic coronary vessel wall cannot be considered inert, but still proves to be a well-adapting ‘organ’.

The ΔRlu course is entirely dependent on the ΔVWd course, but their relationship is only consistent within each triplet and, as such, variable between different triplets. Consequently, the ΔVWd has a dissimilar course within each singular triplet, indicating that the quest for a specific coronary vessel wall dimension as parameter of ‘physiological’ or ‘pathological’ is futile. The same is valid for the intima, media, and adventitia dimensions. Both course-determining factors prove the existence of a strict control mechanism active within all sections and un-altered by the atherosclerotic process.

Successive events describe a system’s course, which is artificial in regards to non-consecutive triplets within each section. This added to the finding that consecutive sections resemble each other only roughly (not shown), assuming systems processes are active throughout the vessel wall. If and how the control mechanism functions in a longitudinal way should be revealed by further investigation.

Overall, the findings indicate that basic systems and mechanisms are un-altered by the atherosclerotic process, which makes a systemic defect as the cause of atherosclerosis less probable and a superposed agent (e.g., inflammation) more plausible. The constituent parts of the coronary vessel wall (intima, media, adventitia) act as a true functional unit via a counter-balancing system and cannot be considered as representative of the coronary vessel wall pathology. If this is also the case with elastic arteries, such as the carotid arteries, using the intima/media ratio as a parameter of coronary artery pathology could be debatable [8, 9]. The stages observed in every section could instigate reconsideration of the cause of coronary artery remodeling [10].

Atherosclerosis is an umbrella term for a process in which plaques are ultimately formed. The aorta appears more ulcerative (craters) than the coronary arteries and the processes leading to these differences uncertain. Therefore, findings on coronary atherosclerotic processes cannot be considered normative for the aortic processes. Differences could exist between muscular and elastic arteries, or even between different muscular or elastic arteries. This is illustrated by the coronary arteries initially lacking the sub-endothelial (intimal) layer, in contrast to cerebral arteries initially lacking the adventitial layer, making them translucent *in vivo*.

Therefore, our study is only the first step in understanding the process from arterial physiology to pathology. Further investigation of different arteries (single sections and section series) could elucidate similarities and/or differences. The random systemic sampling technique could aid in obtaining results for comparisons.

